# Data-driven inference of Boolean networks from transcriptomes to predict cellular differentiation and reprogramming

**DOI:** 10.1101/2024.10.21.618706

**Authors:** Stéphanie Chevalier, Julia Becker, Yujuan Gui, Vincent Noël, Cui Su, Sascha Jung, Laurence Calzone, Andrei Zinovyev, Antonio del Sol, Jun Pang, Lasse Sinkkonen, Thomas Sauter, Loïc Paulevé

## Abstract

Boolean networks provide robust explainable and predictive models of cellular dynamics, especially for cellular differentiation and fate decision processes. Yet, the construction of such models is extremely challenging, as it requires integrating prior knowledge with experimental observation of transcriptome, potentially relating thousands of genes. We present a general methodology, implemented in the software tool BoNesis, for the qualitative modeling of gene regulation behind the observed state changes from transcriptome data and prior knowledge of the gene regulatory network. BoNesis allows computing ensembles of Boolean networks, where each of them is able to reproduce the modeled differentiation process. We illustrate the scalability and versatility of BoNesis with two applications: the modeling of hematopoiesis from single-cell RNA-Seq data, and modeling the differentiation of bone marrow stromal cells into adipocytes and osteoblasts from bulk RNA-seq time series data. For this later case, we took advantage of ensemble modeling to predict combinations of reprogramming factors for trans-differentiation that are robust to model uncertainties due to variations in experimental replicates and choice of binarization method. Moreover, we performed an in silico assessment of the fidelity and efficiency of the reprogramming, and conducted preliminary experimental validation.

## INTRODUCTION

Mathematical models have demonstrated their utility in elucidating experimental findings that might challenge intuitive comprehension. Through meticulous depiction of interactions within intricate signaling pathways and by contextualizing the dynamics of gene expression, these models provide a systematic approach to unveil the regulatory mechanisms that control cellular processes and their dysregulation in diseases. Among the various mathematical formalisms, the Boolean network (Boolean network) is a simple but expressive formalism that relies on pragmatic rules to qualitatively simulate essential systems’ features. It is notably valuable in poorly understood large-scale systems as it can be employed for systems with hundreds of components and as the inference of Boolean network models, contrary to quantitative models (typically ordinary differential equation (ODE)-based models), does not require kinetic parameters derived from in-depth and often unavailable knowledge.

Consequently, Boolean networks are increasingly used to capture the interaction dynamics within complex signaling pathways and regulatory mechanisms governing cellular behaviors. They have been inferred from high-throughput data for modeling a range of biologically meaningful phenomena such as the mammalian cell cycle (19), cell differentiation and specifications (53, 57, 16, 30, 36), stress/aging-related cell behaviors (55, 75), cell apoptosis (50) and cancer cell functions (91, 15, 87, 58). Recent endeavors have focused on enhancing the quantitative interpretation of the resulting models, incorporating probabilistic approaches to effectively simulate heterogeneous cell populations and dynamical interacting populations (74, 76), and introducing a semantics that offers the formal guarantee of completely capturing any behavior achievable by any quantitative model (multilevel or ODE) following the same logic (69).

A Boolean network consists of logical rules that are associated to each variable and that predict how their state evolves with time. The regulators of a variable are combined with logical connectors *or, and*, and *not* and define conditions for a variable to be at 0 (false/inactive/absent) or (1/true/active/present). Through structural and dynamic analyses along with simulations under perturbations, Boolean networks provide versatile opportunities for exploring mechanisms underlying biological phenomena. This versatility allows them to serve as standalone informative tools but also, for specific modeling contexts and needs such as in systems pharmacology, to lay the groundwork for more detailed pharmacokinetic/pharmacodynamic and quantitative systems pharmacology (QSP) models using ODEs (38, 13, 6, 70).

One the prominent challenge for applications of Boolean networks in biology is the design of their logical rules.Indeed, besides the inference of the underlying gene regulatory network, the search for logical rules faces a double combinatorial explosion: the number of possible logical rules for a single variable is exponential with the number of its regulators, and checking whether a candidate Boolean network possesses the desired dynamical features (steady states, trajectories, …) can involve analysis that take time and memory exponential with the number of variables. Thus, in most applications of the literature, the Boolean networks have been manually designed from expert knowledge and by re-utilizing previously published models. Nevertheless, there has been recent progress on the inference of Boolean models from data (81, 67, 60, 25, 68, 24, 85, 1). These methods address the inference problem with different restrictions, either on the type of data and their interpretation in order to obtain simpler dynamical properties, or on restricting the set of logical rules to those having a particular structure, in order to reduce the combinatorics of candidate models. However, these methods remain difficult to scale above hundreds of variables, and typically enforces rather specific way of interpreting the experimental data in terms of Boolean properties.

In this paper, we present a general methodology for the inference of Boolean networks from knowledge, data, and expert interpretation of data, and the computation of prediction from the resulting models. The methodology, summarized in figure 1, builds on the following steps:

**Figure 1:**
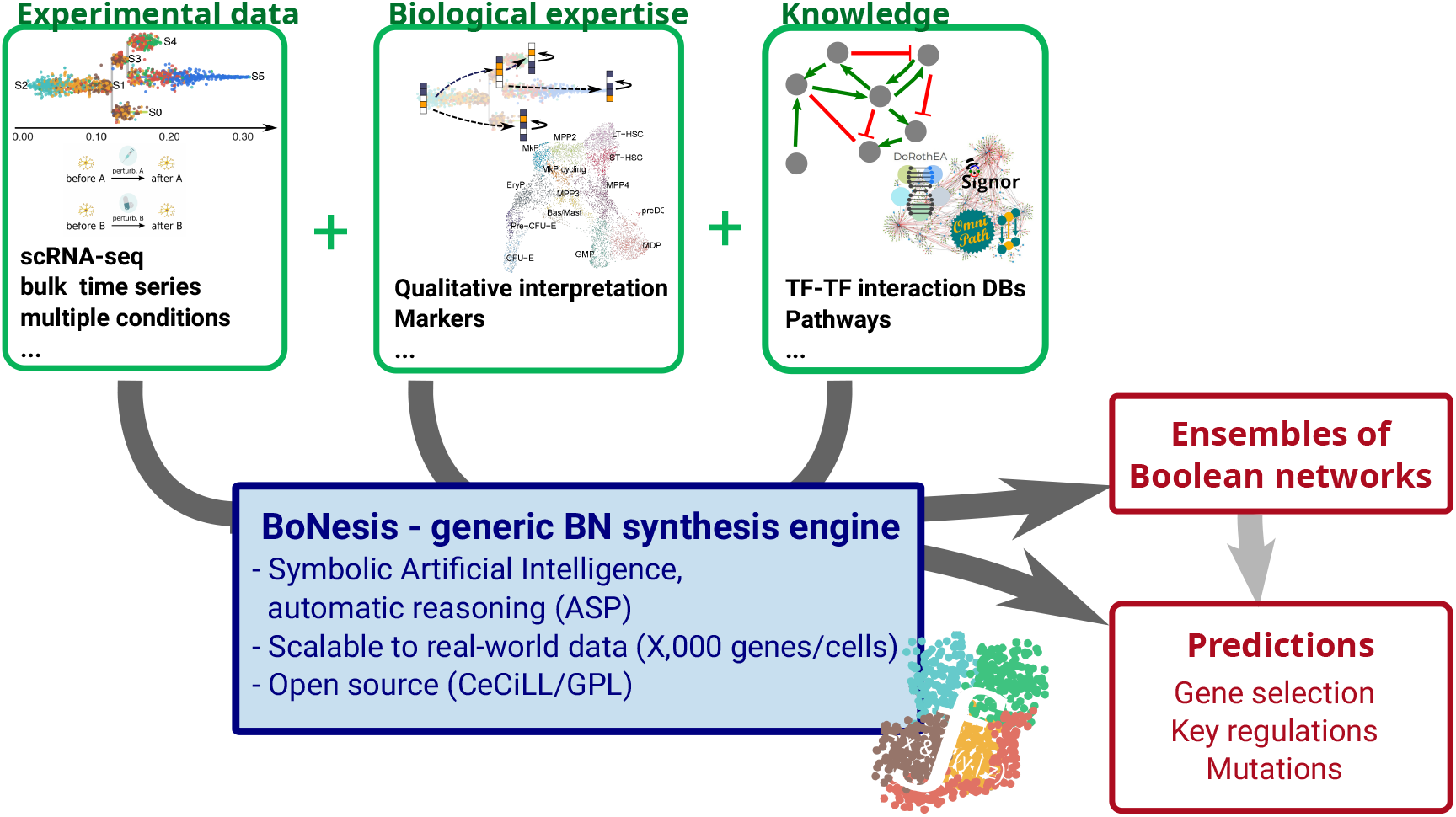
General principle of the conducted inference of ensembles of Boolean networks. By offering a generic modeling language, BoNesis enables integrating prior knowledge on regulation mechanisms with different type of experimental data, after qualitative interpretation, which may depend on biological hypothesis and experimental system. These inputs specify what an admissible model is. Then, employing logic programming, BoNesis can generate ensembles of Boolean networks that fulfill the structural and dynamical properties and, by combining with combinatorial optimization technologies, enables to predict important genes and derive predictions for attractor reprogramming.

1. The modeling of the knowledge, essentially in terms of admissible structure for the models.
2. The qualitative modeling of the data in terms of expected dynamical properties of the model. This steps depends on the biological expertise of the system, and relies on data analysis, notably to classify gene expression into binary values.
3. The tool BoNesis (12), which integrates (1) and (2) and, using logic programming and combinatorial optimization algorithms, infers ensembles of Boolean networks that are compatible with modeled static and dynamical properties.
4. The analysis of sampled ensembles of models to perform predictions, including key genes and reprogramming mutations.

The modeling steps, that define the inference problem, allow a versatile pipeline, that is not tied to specific type of data, or a specific interpretation of them. In some sense, our approach aims at moving the modeling effort from the design of Boolean rules to the specification of the expected features of the model. Then, we employ symbolic artificial intelligence technologies to automatically construct models that satisfy the desired properties.

We illustrate the applicability of the pipeline on two extensive case studies: the inference of Boolean networks from scRNA-seq data of hematopoiesis, with the identification of key genes and the analysis of families of candidate models; and the prediction of reprogramming targets for adipocyte to osteoblast conversion from bulk RNA-seq time series data.

Importantly, these case studies demonstrate that our approach is scalable to TF-scale networks, by starting from complete TF-TF regulatory networks and, through the inference of Boolean networks, are able to automatically identify sub-networks that can explain the observed dynamics.

## RESULTS

### Case study 1: Ensemble modeling of hematopoiesis from scRNA-seq data

We applied our inference pipeline (figure 2A) to the identification of key genes and Boolean rules that can explain the hematopoiesis observed in a mouse sample using scRNA-seq data from Nestorowa et al. (62). Employing trajectory reconstruction and binarization methods, we derived a logical specification of the differentiation dynamics. Then, using BoNesis, we considered any Boolean network employing TF regulations referenced in the DoRothEA database, and automatically identified the sparsest among them that are able to reproduce the differentiation dynamics. We compared the selected genes of importance with an export model of the literature, showing a substantial overlap. Finally, we highlighted the advantage of the ensemble modeling by analyzing the variability of Boolean models compatible with the input data. We notably performed clustering of sampled models, resulting in clear 3 sub-families of models that can be distinguished on specific features of Boolean rules.

**Figure 2:**
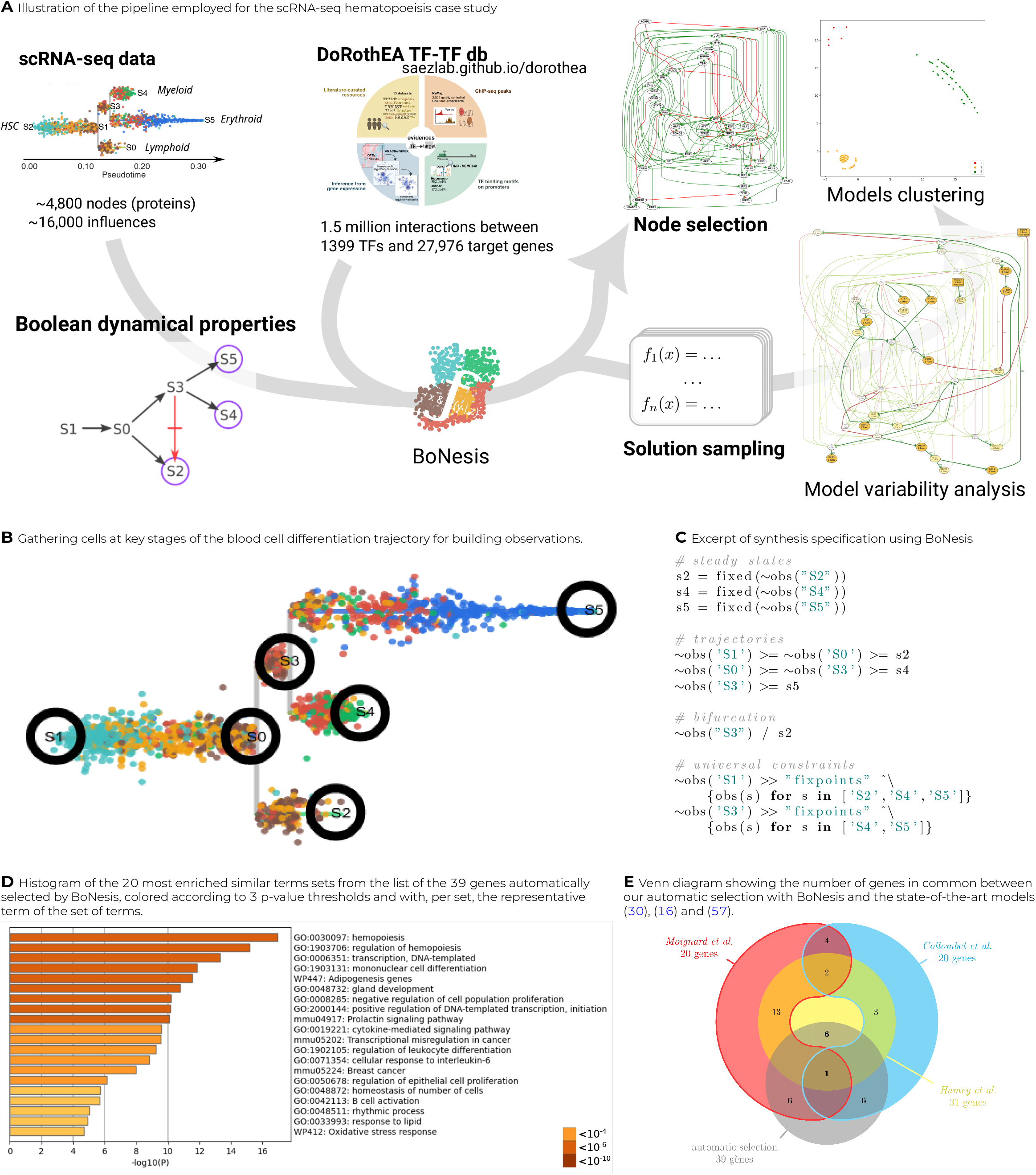
Overview of the case study on scRNA-seq data driven modeling of hematopoeisis.

#### Biological context and experimental data

Hematopiesis is a crucial differentiation process of blood cells for immune system regeneration. It has been extensively studied, including with mathematical and logic dynamical models (30, 16, 57, 32).

We focused on data-driven modeling of the early differentiation of mouse hematopoietic stem cells (HSCs) from scRNA-seq data of Nestorowa et al. (62). The data shows heterogenity of cells during HSCs differentiation, including lympho-myeloid primed progenitors (LMPPs) and common myeloid progenitors (CMPs), further differentiated into granulocyte-monocyte progenitors (GMPs) and megakaryocyte-erythrocyte progenitors (MEPs). We performed hyper-variable gene selection and trajectory reconstruction using Stream (8). The resulting trajectory has the shape of a tree with two bifurcations visible in figure 2B, having as root the endpoint that concentrates the hematopoietic stem cells.

#### Qualitative interpretation of the data

To transform obtained trajectories into properties over Boolean states, we considered *six states* that must correspond to the start and end of branches (named S0 to S5 in figure 2B). Moreover, in order to reduce the sensitivity bias of single-cell observations, we choose to consider observations formed by the union of several cells. This resulted in six clusters of a few tens to hundred of cells, corresponding to initiation (root), two bifurcations points, and three leaves, we considered to be steady states of the Boolean model. We classified the activity of each gene of each cluster using PROFILE (7) on individual cells and an aggregation by majority of value among 0, 1, and ND (not determined).

Then, we specified the expected dynamical properties of a Boolean network corresponding to the data as follows. There must exist trajectories linking with states following the STREAM trajectories: e.g., there must exists a trajectory from a Boolean state corresponding to the root S1 to the Boolean state matching with S0, then from that later Boolean state to a Boolean state matching with S2, and so forth. Moreover, we requested that the Boolean states corresponding to the leaves (S2, S4, S6) must be steady state of the Boolean model, and that any steady state reachable from S3 must match with S5 and S4, and any steady state reachable from the root state must match with S2, S4, and S5. Figure 2C summarizes the declared properties.

#### Gene selection with BoNesis and DoRothEA

Besides the expected dynamical properties of the model, BoNesis requires an *a priori* GRN from which it will reconstruct Boolean rules that (1) employ only genes and regulations referenced in the input GRN, and (2) form a Boolean network that possess the expressed dynamical properties. Moreover, BoNesis is able to identify the genes that can have a constant binary expression towards the whole dynamics, and can thus be ignored in the final model. In the scope of this case study, we took as a priori GRN the full DoRothEA TF regulation database (34) with regulation p to confidence level C, comprising 2,777 regulations among 1,001 genes, 849 of which have an expression measurement in our dataset. We performed a multi-stage combinatorial optimization procedure to identify the largest number of TF genes that cannot be considered as constant, and for each there exists a Boolean network having the desired dynamical properties. It resulted in the selection of 39 TF genes and 137 TF-TF interactions shown in figure S1. A gene set enrichment analysis is performed with Metascape (92) shows a clear enrichment of terms related to hematopoiesis (figure 2D), with the top term being hematopoiesis, followed by others representing more specific biological processes included in hematopoiesis.

Moreover, we compared the data-driven selected genes with three expert art logical models of hematopoiesis by Hamey et al. (30), Collombet et al. (16) and Moignard et al. (57). These three models, composed of 20 to 31 genes, include a total of 53 genes and it is worth noticing that there is no consensus on the interactions implied in the regulation of this process since only 2 genes are common to these three models. The Venn diagram of figure 2E shows the intersection between the components we have automatically selected thanks to BoNesis and the three state-of-the-art hematopoiesis models. Each of state-of-the-art model shares 6 genes with our selection (for a total of 10 distinct genes in common):

- With Hamey et al. (30): FLI1, GATA1, GFI1B, IKZF1, MYB, RUNX1;
- with Moignard et al. (57): FLI1, GATA1, GFI1B, IKZF1, MYB, SPI1;
- with Collombet et al. (16): CEBPA, EFB1, IKZF1, MEF2C, RUNX1, SPI1.

#### Ensemble analysis of compatible models

From the TF-TF subnetwork extracted in the previous step (figure S1), we employed BoNesis to sample 1,000 distinct Boolean networks that all respect the qualitative dynamical properties described previously. The sampling has been performed using heuristics to range over models with diverse logical rules. Each of the sampled Boolean network is able to reproduce the qualitative differentiation dynamics, Moreover, the universal constraints on the reachable steady states, ensure by design, that all trajectories from the root state end in one of the three observed differentiated type. Furthermore, we verified a posteriori that none of the sampled network possess cyclic attractors.

##### Variability analysis shows highly preserved logic rules for two thirds of selected genes

For each selected gene, we analyzed how many Boolean functions has been assigned to it in the sampled ensemble of 1,000 Boolean networks (figure S2). It resulted that 12 genes always received the same Boolean function, 12 other genes received only 2 to 3 distinct Boolean functions. This suggest that a large part of the logic rules are highly preserved in all compatible models. Most of the diversity in the sampled ensembles is essentially focused on gene FOS (682 different Boolean functions) and TRP53 (44 Boolean functions).

##### Clustering identifies 3 sub-families of compatible Boolean models

Clustering the models according to the similarity of their functions can highlight different possible pathways for a process regulation and also point some biologically irrelevant groups. We performed a multidimensional scaling clustering (MDS) of sampled Boolean networks, using distance based on the inequality of Boolean functions: given two Boolean networks *f* and *g* of size *n*, 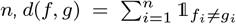. MDS results highlight 3 groups of models (figure S3). It should be kept in mind that the number of models in each group does not reflect their biological relevance, as the group size may result from underlying combinatorial aspects of compatible models, and may also relate to which some close solutions may be easier to find. By comparing the complexity of logical functions across the different cluster, we identified that one of the cluster (the one in red in figure S3C) encompasses Boolean networks with much simpler rules (single activation condition), compared to the two others (Fig. S4-S6).

Thus, the ability to cluster the models and analyze their variability enable to pinpoint model features that can be challenged with expert knowledge are further experimental studies in order to discriminate among sub-families of models.

### Case study 2: Prediction of reprogramming targets for adipocyte to osteoblast conversion from bulk RNA-seq time series data

This case study demonstrates a full pipeline going from experimental bulk RNA-seq time series data and background knowledge on TF-TF networks to prediction of genetic mutations for trans-differentiation and preliminary experimental validation. The predictions have been obtained by combining inference of Boolean networks ensembles with formal methods for control and ensemble simulations for scoring (figure 3).

**Figure 3:**
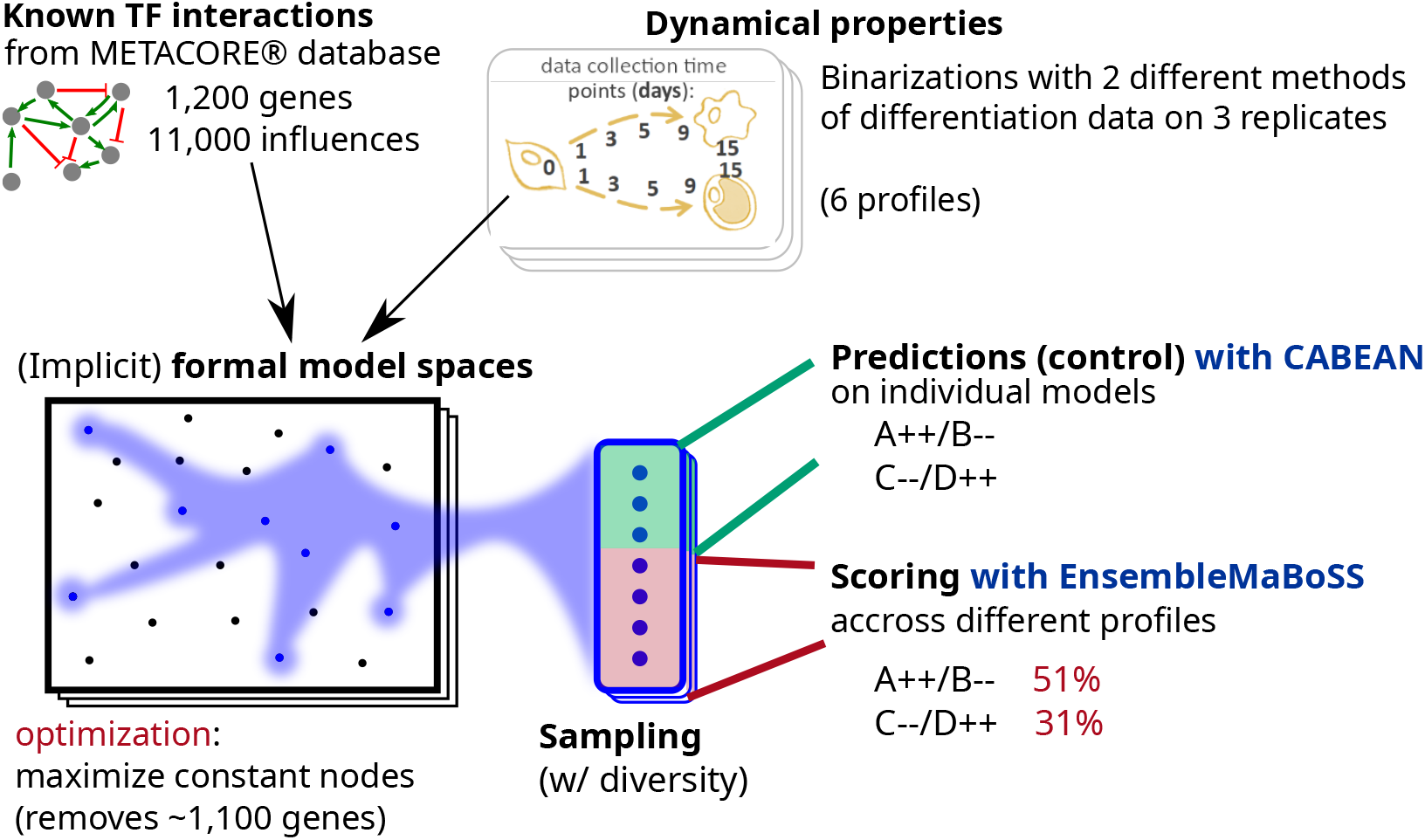
Illustration of the pipeline employed for case study 2. We combined prior knowledge TF-TF interactions extracted from METACORE(r) database with variants of Boolean dynamical properties extracted from time series bulk RNA-seq data. We employed BoNesis to sample Boolean networks fulfilling these properties. Then, we performed prediction of reprogramming determinants using CABEAN on a subset of these sampled models. The identified reprogramming determinants are combinations of gene knock-outs and constitute activations. The predictions are then assessed on the other sampled Boolean models using EnsembleMaBoSS simulation, leading to a scoring in terms of fidelity and efficiency, and thus a ranking of most robust predictions.

#### Biological context and experimental data

Bone marrow stromal progenitor cells (MSCs) are multipotent cells capable of differentiating either into osteoblasts to form bone tissue or into bone marrow adipocytes, that play an important role in the hormonal homeostasis of the bone marrow (2). These cells are reciprocal in their differentiation and correct balance is important for bone health, with increased adipogenesis observed in obesity and during aging. Previous studies have sought to understand the gene regulatory networks underlying MSC differentiation and improved understanding of these networks could allow for identification of efficient reprogramming targets and better control of bone marrow cell composition (54). We have previously performed transcriptomic profiling of ST2 cells, a mouse MSC cell line, across multiple time points of parallel differentiations into adipocytes and osteoblasts using RNA-seq (23). Both differentiations were performed for 15 days with RNA samples collected from undifferentiated cells (ST2D0) and at 5 different time points (D1, D3, D5, D9, and D15) of each differentiation. Osteoblastogenesis (OD1-OD15) was performed with one composition of differentiation medium for the entire duration of 15 days while for adipogenesis (AD1-AD15) two different media were used, with media composition changed on the third day of differentiation (AD3). *In vitro* differentiations are known to vary between experiments (26) and therefore three independent differentiation experiments were performed to capture the robustly reproducible gene expression changes. Please see Materials and Methods for further details. The obtained dataset formed a rich resource of dynamic gene expression profiles towards two different trajectories from the same starting point.

#### Qualitative modeling of bulk RNA-seq time courses triplicates

##### Bulk RNA-seq binarization

We employed two different binarization methods by classifying gene activity with respect to background RNA-seq data on a range of tissues, either by bootstrapping parametric distributions (RefBool), or by a simpler statistical procedure applying gene-specific cutoffs, referred to as MUQ in the following. Each gene of each time point is thus assigned to a Boolean or undefined value for each binarization method and each replicate. In total, 1560 genes received a binary value in at least one binarization method and one replicate. For a fixed binarization method, we observed opposite classifications between replicates for up to 29 genes, while up to only 2 genes when comparing across binarization methods. This can be explained by the fact that, in general, RefBool classifies less genes than MUQ, and genes classified by RefBool are classified by MUQ similarly. In order to account for this variance in binarizations, we considered each of the 6 profiles (2 binarization methods times 3 replicates) as alternate model specifications. Our rational was to constitute ensembles of models for each of these profiles, and study the robustness of reprogramming prediction across them.

##### Dynamical properties

Our main modeling hypothesis was that the Boolean model must be able to reproduce the observed maturation trajectories from fixed cellular environments. For instance, due to the change of treatment between days 1 and 3 of adipocyte culture, we did not required the existence of a trajectory from `AD1` to `AD3`. From the experimental protocol, this resulted in the specification of two trajectories: the maturation of adipocytes, modeled as the existence of a Boolean trajectory going through `AD3 → AD5 → AD9 → AD15;` and the maturation of osteoblasts, modeled as the existence of a Boolean trajectory going though `OD1 → OD3 → OD5 → OD9 → OD15`.

Moreover, we assumed that ST2D0, AD15, and OD15 are observations of cells in steady states. At last, we modeled the observed cellular differentiation process by denying the existence of trajectories across the two branches, nor reverting to the precursor state: there must not exist a trajectory from `AD3` to `OD15`, from `AD3` to `ST2D0`, from `OD1` to `AD15`, nor from `OD1` to `ST2D0`.

#### Gene selection and sampling of ensemble of diverse Boolean models

We performed selection of genes and sampling of diverse ensembles of Boolean networks from the qualitative modeling of RNAseq data. We followed a workflow similar to use case 1, that we repeated for each replicate and for each binarization method. Then, as the gene selection resulted in several optimal solutions, the sampling of Boolean networks has been repeated on each of them.

The prior GRN was consisting of TF-TF interactions extracted from METACORE database from all the TFs of the RNA-seq dataset. It resulted in a signed digraph comprising 1,027 genes with 11,159 regulations, 169 of which were with an undetermined sign. This prior GRN served to define the set of candidate Boolean networks. Because several genes have more than one hundred referenced regulators, we restricted to the Boolean networks whose activation functions can be expressed with at most 32 disjunctive clauses, without any limit on the size of the clauses. Thus, while we permit a gene to depend on all of its potential regulators, we limited the number of activation contexts.

On the 1,027 genes of the prior GRN, one can expect that only a fraction of them are involved in the observed differentiation process. As in case study 1, we employed BoNesis to perform gene selection by identifying Boolean networks reproducing the dynamics of as much as varying gene as possible, while assigning as much as activation function as possible to a constant value. Depending on the replicate and binarization method, the optimization results in different optimal sets of genes to preserve, ranging from 49 to 79 genes. Finally, we performed the diverse sampling of 264 distinct Boolean networks for each of the six profiles, resulting in 1,584 Boolean networks verifying the qualitative dynamical properties corresponding to at least one replicate and one binarization method, and using one of the optimal set of genes.

#### Prediction of reprogramming targets with high fidelity and efficiency

Our objective was to predict combinations of perturbations of gene expression to trigger a transdifferentiation of adiocytes into osteoblasts. In terms of Boolean network, this corresponded to identifying control strategies to enforce the reachability of the `OD15` state from the `AD15` state. In order to account for candidate model heterogenity, our approach was to compute reprogramming targets on individual Boolean networks from a subset of the sampled ensemble, and evaluate them on the full ensemble. In the end, we aimed at selecting the perturbations predicted to be most effective on a range of models reconstructed from different binarization and replicates.

##### Reprogramming computation from subsampled individual models with CABEAN

We selected the CABEAN tool (77) to compute combinations of temporary gene knock-out and constitutive activations enforcing the convergence to `OD15` state from `AD15` state for a given individual Boolean network. For each of the 6 qualitative profiles, we applied CABEAN on 24 of the 264 Boolean networks sampled in the inference part. CABEAN failed on 15 of these 144 Boolean networks due to memory issue. On the remaining 129 models, CABEAN identified combinations of up to 5 simultaneous perturbations leading to a reprogramming from `AD15` to OD15. Because we are interested in perturbations that can be effective on as many models as possible, we kept combinations of perturbations that have been identified in at least 10% of the individual models given to CABEAN. This short list of candidates contained 34 different combinations of 2 to 4 simultaneous perturbations.

##### Simulation of reprogramming with EnsembleMaBoSS

The reprogramming perturbations computed by CABEAN are guaranteed to be effective on their input individual Boolean network. Our objective was to assess the robustness of these (combination of) perturbations the Boolean network ensembles inferred from different qualitative interpretation of the data. To do so, we extended the MaBoSS stochastic Boolean network simulator to sample trajectories from the Boolean network ensembles: each Boolean network of the ensemble is simulated *k* times from the corresponding the state corresponding to `AD15` while enforcing the given reprogramming perturbation. For each candidate combination of perturbations, we obtain an estimation of the distribution of the steady states of Boolean networks of the ensembles after enforcing the perturbation from the `AD15` state.

##### Evaluation of reprogramming efficiency and fidelity

We defined scores to evaluate reprogramming candidates from their simulation on Boolean network ensembles for their ability to reprogram to the osteoblast phenotype, as observed at OD15. Inspired by usual cellular reprogramming assessments, we considered two measures: the *efficiency* relates to the proportion of cells (models) that show all the prior knowledge adipocyte gene markers (ADIPOQ, FABP4, CEBPA, LPL) and none of the osteoblast prior knowledge marker gene (ALPL, HEY1, SP7). Then, the *fidelity* relates to the similarity of those cells to the full `OD15` state.

For a given reprogramming perturbation, we write *S* the set of states resulting from the EnsembleMaBoSS simulation, and for each state *s* ∈ *S*, we denote by *p*_*s*_ its estimated steady state probability. Moreover, we define **1**_Ost_(*s*) as being equal to 1 whenever each osteoblast marker gene is active in *s* and each adipocyte marker gene is inactive is *s*, otherwise, **1**_Ost_(*s*) is equal to 0.

The reprogramming efficiency is computed from the estimated steady state distribution after perturbation of Boolean network ensembles as the fraction of simulations ending in a state having all the osteoblast marker genes active and all the adipocyte marker genes inactive:

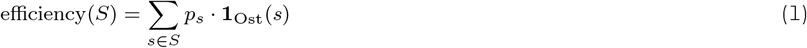

The repgrogramming fidelity employs a similarity measure between a full Boolean state *s* ∈ *S* and the binarized state *β*_OD15_ of the corresponding qualitative interpretation. Because the binary sate of some genes of *β*_OD15_ can be non-determined, the similarity is

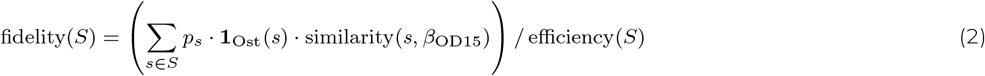

defined as the proportion of binarized genes in *β*_OD15_ that have the same value in *s*. In the end, the fidelity is the expected similarity of the Osteoblast steady states of Boolean networks in the ensemble after reprogramming from AD15:

#### Validation of node selection and reprogramming targets with literature

The TFs included in the predicted reprogramming determinants were ranked by frequency and maximal predicted efficiency (figure 4). Analysis of the literature related to the predicted top TFs revealed their existing association with regulation of osteoblastogenesis, thereby providing indirect validation for our approach. For example, HEY1, a well-known target of Notch signaling(83) and a regulator of osteogenesis(72) was among the factors with highest predicted efficiency score. Similarly, CEBPA, SP1, and TRP63 have been associated with adipogenesis, osteogenesis, and bone formation (45, 63, 27), respectively. With repression of CEBPA likely to act to reverse adipogenic gene expression program. Likewise, the repression of NR2F2 or GATA3 were predicted to lead to high efficiency of conversion, something that is also supported by literature (88, 40, 42, 86). One of the top TFs included TCF7L2, a mediator of WNT signaling (figure 4). Consistently, WNT signaling is known to positively regulate osteoblastogenesis (17, 3), with the effect mediated by TCF7L2 (90). Finally, also MYC genes have been previously shown to contribute to reprogramming of fibroblasts towards osteoblasts (89, 56). However, some of the well established regulators of osteoblastogenesis, such as SP7 (also known as Osterix) or RUNX2 (89, 56), were not included among the predicted reprogramming determinants. This could be related to, at least in case of RUNX2, its already abundant expression in undifferentiated ST2 cells (23).

**Figure 4:**
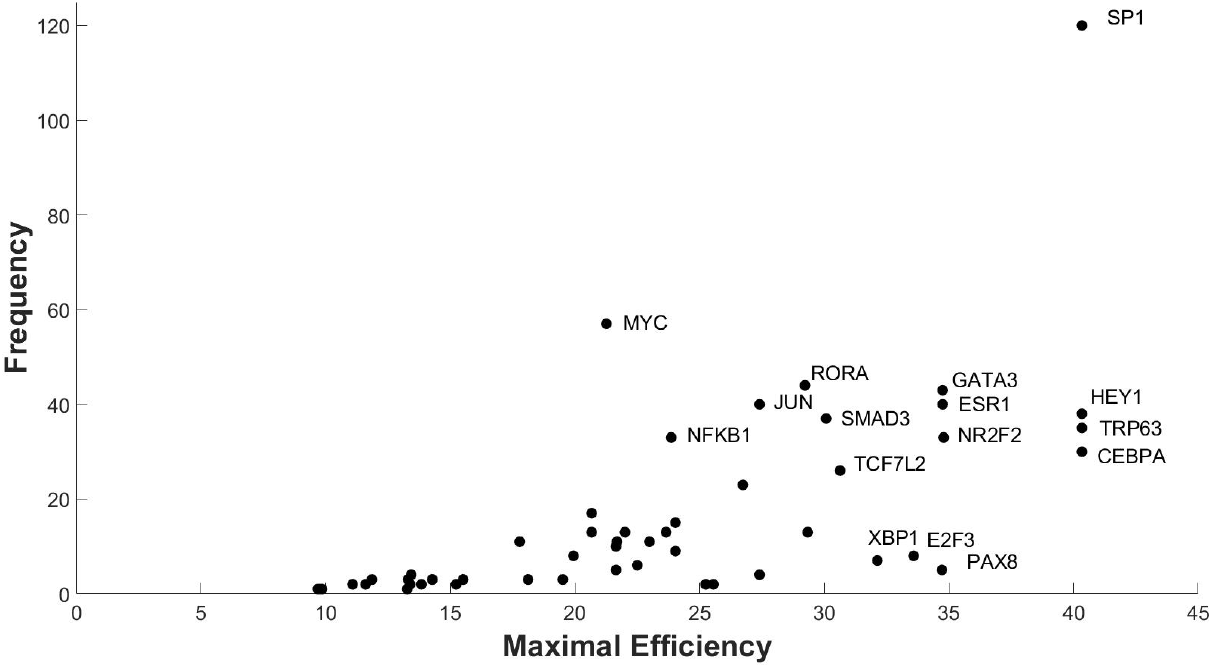
Specific TFs occur more often and with higher efficiency in the predicted reprogramming determinants. Frequency (y-axis) and maximal efficiency (x-axis) of the TFs included in the predicted reprogramming determinants applying permanent stimulation are reported. Names of most promising TFs are given.

Based on the predicted combinatorial efficiences, and the existing literature evidence, three combinations of three determinants each were selected for experimental validation. Reprogramming experiment 1 (*RE1*) involved up-regulation of TRP63 and downregulation of CEBPA and SP1. *RE2* involved up-regulation of TCF7L2 and TRP63 and down-regulation of SP1. And *RE3* involved up-regulation of TCF7L2 and down-regulation of CEBPA and SP1.

#### Preliminary experimental validation

In order to experimentally test the different combinations of reprogramming determinants selected for *RE1, RE2* and *RE3*, we took advantage of lentiviral overexpression in combination with siRNA-mediated gene repression. Overexpression constructs were synthesized at Sirion Biotech and were designed to coexpress either green fluorescent protein (GFP) and red fluorescent protein (RFP) to confirm a successful lentiviral transduction and to allow to separate between different constructs (specifically co-transduction of TCF7L2 and TRP63). Day 9 adipocytes were transduced with lentivirus of interest and 24 h later transfected with the relevant siRNAs to achieve the respective combinations for each RE. In parallel control cells were transduced with a negative control virus overexpressing only GFP and subsequently transfected with a scrambled control siRNA (please see Materials and Methods for additional details). Using reverse transcription real-time quantitative PCR (RT-qPCR) TCF7L2 and TRP63 were confirmed to be 8- and 15-fold overexpressed in undifferentiated cells, respectively, while siRNAs against SP1 and CEBPA led to 35-68% decrease in their mRNA expression (figure S7). Following transfections, the adipocytes were cultured in osteoblast medium for 9 days before collection of the cells for single cell RNA-sequencing (scRNA-seq). After the sequencing and data processing, transduced cells were identified based on the presence of the mRNA sequence encoding for GFP and/or RFP and the relative changes in the transcriptomes were analysed. Differentially expressed genes (DEGs) were obtained in the single-cell expression data based on the Wilcoxon rank-sum test with FDR<0.05. Directionality of the observed change was determined with the common language effect size, with a >0.5 indicating up-regulation (see Supplementary Table 1). The precision for the upregulated single-cell DEGs compared against the bulk DEGs (Day 15: Adipocyte vs Osteoblast, adjusted p-value<0.05) is 0.57, 0.51, and 0.33 for the three different reprogramming experiments *RE1-RE3*. This means that for *RE1* 57% of the genes which were called up-regulated in the single-cell experiment were also up-regulated in osteoblasts compared to adipocytes in the bulk data (23), indicating an initial consistent effect of the applied reprogramming determinant. The respective hypergeometric p-values for enrichment of the true positives in the single-cell DEGs are 4.4e−14, 1.1e−12, and 0.057, respectively. This promising tendency is also confirmed by visual inspection of the bulk RNA-seq data of DEGs obtained in the single-cell experiment (for *RE1* see figures S8 and S9). The very low recall (*<* 0.02)in all experiments however indicates that only few of the relevant genes responded to this perturbation in the relatively short time frame of the experiment.

## DISCUSSION

### Overview on the methodology

We demonstrated our methodology on two complementary case studies for a combination of data-driven and expert-driven inference of Boolean models from scRNA-seq and bulk RNA-seq data. The essence of our approach is the explicit modeling of the inference problem, with the specification of the prior knowledge, typically extracted from databases of TF-TF interactions, and the specification of the expected properties of the model. These latter must reflect the expert and qualitative interpretation of the experimental data. We relied on the BoNesis engine which enables to specify and combine a broad range of dynamical properties, notably related to trajectories and steady states. Then, ensembles of compatible models can be sampled and further analyzed. We took advantage of ensemble modeling to analyze candidate model variability with model clustering (case study 1), as well as to provide robust reprogramming predictions (case study 2).

The flexibility of the workflow facilitates the comparison of different modeling choices and hypotheses. For instance, in case study 2, we considered different binarization methods and different replicates. We took advantage of the variability to generate ensemble of models spanning the different hypotheses and identify consensus predictions. The choice of the input GRN can also impact the results, and it could be assessed in a similar fashion. Such an analysis is out of the scope of this paper, as our objective was not in the benchmark of prior knowledge data nor binarization methods.

Remarkably, the inference pipeline was tractable on TF-scale networks. By employing logic programming and relying on recent complexity breakthrough in Boolean network analysis (69), we have been able to fully account for thousands of transcription factors. Then, we took advantage of combinatorial optimization technologies to automatically prune non-necessary variables and identify sub-networks that drive the observed dynamics.

#### Case study 1

With this case study, we leveraged ensemble modeling to analyze diversity of models that can explain the observed differentiation process. The experimental scRNA-seq data has been processed with trajectory inference methods from which we extracted both clusters of cells corresponding to initiation, bifurcation, and differentiated states, and dynamical properties related to existence of trajectories linking those states.

From a full TF-TF interaction database, we have been able to identify a subset of core TFs from which can be drawn Boolean networks reproducing the qualitative differentiation, that present some intersection with expert models from the literature. By analyzing the ensemble of sampled models, we identified three families of models that diverge by the complexity of their logical rules The variability analysis also emphasized model patterns that are preserved across the ensemble.

As we performed a partial enumeration, we are not assured not to miss models with characteristics different from those of the three highlighted groups. However, partial enumerations of 250 and 500 models already highlight the three groups and suggest that increasing the number of models only leads to an increase of the size of the 3 groups, without highlighting any new type of models. This motivates the chosen number of 1000 models. MDS done with 250 and 500 models are presented in figure S3A and figure S3B.

This modeling of hematopoiesis from a causal network automatically inferred from the dynamics of the data already provides avenues for further exploration of the mechanisms of its regulation, with the possibility of ensemble simulation of models with MaBoSS (76).

#### Case study 2

The applied pipeline yielded promising predictions for the reprogramming of adipocytes to osteoblasts, as confirmed by our literature validation, highlighting its utility in identifying novel targets from time-series RNA-seq data. Notably, transcription factors such as SP1, HEY1, and TRP63 frequently appeared in both temporary and permanent perturbation lists, suggesting central roles for them in adipocyte-to-osteoblast conversion. The successful prediction included factors like CEBPA and SP1, known to be crucial for adipocyte and osteoblast functions as modulators of the activity of the respective master regulators of these lineages, PPARG and SP7 (also known as Osterix) (49, 64). These TFs’ selection as reprogramming targets is consistent with their involvement in these regulatory networks, supported by their high occurrence in prediction lists and established biological roles in these processes (46, 5).

The reliability of the predictive model is evidenced by its consistency with existing literature on differentiation of mesenchymal cells. Factors such as ATF4 and NR2F2, previously implicated in positive and negative control of osteoblast differentiation, respectively, are found among our predictions (51, 41). Similarly, the positive effect of HEY1 and TRP63 on bone formation in the literature (73, 28) highlights the utility of our approach. The predictions also drew attention to the role of less commonly studied factors like AHR and RARA, which are implicated in sensing the cellular microenvironment crucial for differentiation. AHR, predicted for upregulation in two combinations, is known to shift the gene regulatory network towards a less specialized cell state, potentially facilitating the reprogramming process (29). Similarly, RARA, noted for its negative regulatory role in adipogenesis, is part of the reprogramming predictions (59), suggesting that its modulation could inhibit adipocyte traits while promoting osteoblast characteristics. Follow-up studies of our findings could benefit the understanding of bone-related diseases like osteoporosis. By utilizing knowledge of specific gene targets that promote osteoblast differentiation, new therapeutic avenues may be explored that enhance bone regeneration and repair (66, 14). Additionally, manipulating adipocyte-to-osteoblast conversion could benefit obesity-related diseases where abnormal adipogenesis is prevalent (9).

Nevertheless, for now our validation experiments do face limitations. While gene expression changes upon reprogramming perturbations were enriched for genes involved in osteoblastogenesis, as observed by scRNA-seq, the overall efficiency of gene delivery and knockdown would need to be improved to be able to better address the quality of our predictions. Usage of stable and inducible systems that would be independent of chemicals such as doxycycline, that might perturb cellular systems beyond intended effects, should optimally be used. Additionally, the complexity of gene and protein interactions, including the necessity of considering interaction partners like beta-catenin with TCF7L2 or post-translational modifications like phosphorylation of factors like JUN and FOS, suggests a greater number of (also non-genetic) perturbations may be required for effective reprogramming than initially predicted. Integrating multi-omics data with transcriptomic profiles could provide a more comprehensive understanding of the reprogramming processes, allowing for the examination of post-translational modifications and protein activity states crucial in the transcriptional state changes.

## MATERIALS AND METHODS

### Boolean network inference methods

#### Knowledge-based (bottom-up), data-based (top-down), and combinations

The manual approach is a bottom-up modeling (also called forward modeling) that designs a model through expert and literature-based knowledge to determine the Boolean functions from known and suspected interactions (19, 82, 91, 50, 53, 15, 16, 55, 75, 36, 87). The resulting model is validated or refined in an iterative process according to the fitting between its dynamics features and observations of the biological phenomenon (for example its attractors correspond to known phenotypes). This time-consuming approach requires a deep understanding of the biological system and does not ensure that all possible regulations leading to the observed behavior are explored. In contrast, methods have been developed to propose a top-down (reverse) modeling approach that derives from experimental data both the topology of the network, namely the causalities between the biological components, and the logical rules of nodes activation, constituting the Boolean network (18, 57, 84, 30, 47, 35, 71, 20, 33). This purely data-based inference approach can suffer from overfitting to a dataset and circular reasoning: a dataset generates a network, which then predicts the dataset. It also confronts the vastness of the solution space without considering prior knowledge about the phenomena. Consequently, other methods have been developed midway between these previously mentioned top-down and bottom-up modeling approaches (81, 67, 60, 25, 68, 10, 24, 85, 1), which encompasses the method presented here. The purpose is to leverage both experimental observations and prior knowledge on interactions related to the biological phenomenon to model. Each tool combining data- and knowledge-based modeling considers as input both experimental data, typically expression profiles, and a gene regulatory network (GRN) potentially extended with some features. The GRN, also named influence graph or regulation graph, structures the prior knowledge about interacting components according to the following definition: a directed graph that represents regulatory interactions between biological components, interactions categorized in the simplest case as activations and inhibitions (52). A high number of tools have been developed to address these issues because they perform the Boolean network inference through different algorithms and are designed for different types of experimental data and different biological interpretations of it.

#### Partial vs exhaustive exploration of the space of solutions

Exploring the space of possible Boolean networks and their dynamics is a challenge because there can be millions of combinations of logical rules that can be formulated. This is why, depending on the algorithm implemented by the tool, the output models can be either Boolean networks that exactly reproduce the desired properties, or Boolean networks optimized to best match these properties with no guarantee of optimality. In the first case, the exploration of solutions can be exhaustive. It should also be taken into account that this search for models (exact or by optimization) can be limited to a subspace of solutions if Boolean function patterns are imposed. ***Optimization problems vs satisfiability problems***. If we focus on the way solutions are explored, we can distinguish between optimization-based methods (44, 68, 47, 81, 1, 35, 20) (heuristic, identify a Boolean network that best aligns with the observations and prior knowledge, based on a defined set of criteria or objective functions) and decision-based methods on satisfiability problems (67, 30, 10, 1, 85) (exact, determine whether there exists a Boolean network consistent with the observations and prior knowledge – based on constraints on its topology and dynamics – and find such network if it exists). The methodology we present in this paper, based on previously published works (10, 11), belongs to this latter category. According to the chosen approach, some methods output (near-)optimal solutions with no guarantee of reaching a global optimum, while others can theoretically explore the solution space exhaustively to output the whole set of Boolean networks that comply exactly with the biological constraints (in practice often limited to a subset of these Boolean networks as the network size increases and the number of solutions becomes tremendous). This latter ability of inferring an ensemble of models is highly informative. It enables variability analysis across the models to identify common patterns of interactions, supporting the formulation of hypotheses that benefit both the modeling process and the biological exploration of key components.

#### Patterns of Boolean functions

Beyond this distinction between categories of algorithms, limitations on the formulation of the Boolean rules that are considered by the methods also impact which solutions are explored. It can be a fixed limit on the number of regulators (e.g., a maximum of 6 regulators including 4 activators and 2 inhibitors (30, 33)) or a certain pattern of rules (e.g., a component is expressed if all its activators are expressed and none of its inhibitors, i.e., *and* logic gate between activators and *or* between inhibitors (18, 33)). Because of the combinatorial explosion, methods designed to consider all possible Boolean functions can also offer a limit on the size of the Boolean function to enable dealing with dozens of components and complex regulations. This limit can be on the number of regulators or, like the choice we made, on the number of different combinations of regulators (i.e. a limit on the number of clauses that can compose the Boolean functions). This latter choice seems to be biologically consistent: the number of direct regulators for a gene or protein can vary significantly, often being numerous. Whereas the number of functional states, reflected by specific combinations of these regulators, is typically more constrained, ensuring system resilience and controlled responses to a manageable range of conditions.

#### Different qualitative interpretation of experimental data

Regardless the implemented algorithm, the Boolean network inference tools can also be judiciously compared by focusing on the types of data that they can consider for the modeling: bulk or single-cell expression profiles, at cellular steady states or covering cellular evolutions, linear evolutions, or with bifurcations (e.g., lineage differentiation), observed with time series limited to two time points or longer, with/out perturbations on components, etc. This is directly linked to the dynamical features they are able to model. ***Steady-state data***. Measurements acquired at assumed stable cellular states are commonly interpreted as attractors in the Boolean network dynamics. On this basis, some inference tools such as CellNOptR (81), Griffin (60) and BONITA-RD (68) are specifically designed for inferring Boolean networks that respect constraints on their attractors (focusing mostly on fixed points), constraints that are derived from a set of steady-states data. ***Time series data***. A range of tools extends the modeling scope by addressing time series of expression measurements, representing a succession of cellular states ensuing one after the other. Among them is CellNOptR-dt (81) which can fit time course data using a synchronous updating scheme. The modeling of this data is widened to asynchronous semantics with ASKEed (85) and IQCELL (33) that model time series as trajectories in which the succession of measurements constitutes a succession of transitions (reachability in a single dynamic step). Caspo-TS (67), also asynchronous, relaxes the constraint of observing each transition as it considers the reachability property between the observed cellular states. It is worth noting that none of these methods takes into account the modeling of bifurcations/branching in the trajectories. ***Single-cell data***. Among the methods specially designed for inferring Boolean networks from single-cell expression data, we can distinguish different interpretations of their dynamical features. SCNS (57), BTR (44) and SgpNet (20) interpret the set of single-cell measurements as a state space reached from an initial state in an asynchronous Boolean network. These tools seek, through optimization, a Boolean network for which the set of states contained in its transition graph (the “model state space”) is close to the set of binarized expression data (the “data state space”). However, the strategy presented in (71) is based on another interpretation: the state of each single-cell measurement is assumed to be a potential predecessor or successor of the state of any other single-cell measurement coming from the same sample. Consequently, a solution is here a synchronous Boolean network whose dynamics matches a random set of two-time-points time series, namely couples of single-cell measurements. The previously mentioned IQCELL (33) proposes a third interpretation of this data, considering a pseudo-time ordering of the cells seen as a time series and searching for an asynchronous Boolean network whose dynamics includes a sequence of states compatible with the cell ordering. ***Complex dynamics***. A tool that can consider a diversity of dynamical features can exploit different types of data and address the modeling of varied and complex cellular behaviors. Three methods stand out for the complexity of the behaviors that can be modeled. The tool BRE:IN (25) allows for more nuanced and complex dynamic behaviors than the previously mentioned tools as it supports complex temporal logic specifications (CTL and LTL), both in synchronous and asynchronous semantics. However, the high computational cost of checking these traces prevents scaling to networks of biological interactions of dozens of components: the larger networks for benchmarks (25) had 16 nodes with synchronous semantics and 11 with asynchronous. Similarly, Boolean network sketches (4) also performs the inference with the help of modelchecking methods, in asynchronous semantics, but with the richer logic HCTL (hybrid extension of the branching-time temporal logic CTL) allowing more expressive specifications. Unlike BRE:IN, the exploration of the Boolean functions is not limited by patterns, and this method was tested in a network with more than 321 nodes.

### Binarization of transcriptome data

#### Case study 1

From the full scRNA-seq dataset, we employed PROFILE (7) to classify in each cell each gene to either 0, 1, or undetermined, by comparing its expression to the distribution in all cells. Then, each cluster identified using STREAM (see main text) has been binarized using a majority rule: a gene is classified following the majority of values the gene has been classified to in the individual cells of the cluster. Among the 4768 hyper-variable genes selected by STREAM, 1519 have been classified with different binary values in at least two different clusters, and 1369 are classified as binary value in all clusters.

#### Case study 2

We previously performed experiments for acquiring RNA sequencing data in populations of ST2 cells at different stages of differentiation towards adipocytes and osteoblasts (23). Adipocyte differentiation was induced using a medium of isobutylmethylxanthine, dexamethasone, and insulin for 2 days, followed by a change of medium with rosiglitazone insulin (Sigma-Aldrich, I9278) until 9 days. Differentiation towards osteoblasts was induced using bone morphogenetic protein-4 until 15 days. Each experiment has been replicated 3 times. At days 0, 1, 3, 5, 9, 15, a subpopulation of the cells has been sequenced genome-wide. In the scope of this project, we focused on the activity of transcription factors (TFs) in the different stages of the ST2 differentiation. To enable building of qualitative models for the differentiation process, an automated method was applied for transforming the quantitative RNA-seq measurements of TF activities into qualitative assessment of their activity: active (1), inactive (0), or undetermined (intermediate). Two different methods of binarization were employed and compared: RefBool (37) for binarization with respect to background RNA-seq data collected in a range of different cell types; and a statistical analysis developed in the scope of the project, for binarization with respect to the collected RNA-seq data. For this statistical analysis, representative background distributions of gene expressions in 63 mouse tissues were generated from ArchS4 (39) data [https://maayanlab.cloud/archs4/, Kallisto raw read counts, retrieved 2019]. Independent vertex sampling was performed per tissue to remove correlated samples. Samples were further filtered for overall read counts between 10 and 100 Mio and a median>=1 to remove unusual distributions and outliers. This background data was merged with our own data (after Kallisto alignment) and then quantile normalized and converted to TPM (gene length normalization and TPM scaling). The gene-specific background distributions were then applied for discretization of the own data as follows: Gene expressed in own data below median of background was discretized to 0 and above upper quartile to 1. In addition, genes with TPM<1 are discretized to 0. Furthermore, genes with large expression differences over all samples and timepoints were identified via k-means-clustering (2 clusters), with a minimum three-fold-change of centroid locations, at least three data points per cluster, and full-filling a ttest2 (MATLAB©) between the two clusters. 36 genes were found accordingly. The upper cluster was discretized to 1 and the lower cluster to 0.

### Prior knowledge on gene regulation

#### Case study 1

We extracted TF-TF and TF-gene interactions referenced in the DoRothEA database (21) with confidence levels A to C. It resulted in a signed directed graph with 5 186 components and 12895 regulations. We filtered the components to keep only genes with classified binary values, reducing to 1001 components and 2777 regulations.

#### Case study 2

We extracted from the METACORE© database (in 2019) all known interactions (Transcription regulation, Influence on expression, and Binding) between TFs (31) and seven known marker genes which show clear expression changes from adipocytes to osteoblasts in our own expression data: Four high in adipocytes (ADIPOQ, FABP4, CEBPA, LPL) and three high in osteoblasts (ALPL, HEY1, SP7). Only measured nodes and their interactions were kept. This resulted in a network of 1,027 nodes (almost only TFs) with 11,000 pairwise interactions.

### Boolean networks

### Inference of Boolean networks from structural and dynamical properties

The strategy behind BoNesis is to describe, in the form of a logical problem to be solved, the search for a Boolean network compatible with a network architecture and with given dynamic properties. To do so, BoNesis integrates prior knowledge influence graph and observations within the same logic program, so that any solution of this program is a Boolean network made up of possible interactions given the PKN whose dynamic properties are compatible with the behavior of the observations.

The logic program is written in Answer-Set Programming. It describes the biological data (the prior influence graph as well as the observations and the dynamical properties linking them) and the modeling formalism (the Boolean network and the computation of its dynamics in Most Permissive Semantics) via predicates and constraints. The constraints are necessary and sufficient conditions that guarantee that any solution of the problem is a Boolean network compatible with the biological data, i.e., a Boolean network included in the architecture of the prior influence graph and whose dynamic properties are compatible with the behavior of the observations. Solutions are obtained using an answer-set solver, *clasp* (22): the models are the answer-sets satisfying the logic program.

#### Component selection

Building a prior knowledge influence graph specific to a biological process is a complex and delicate expert task. Yet, determining which interactions are to be considered when building a model is an essential step in the preparation of data before synthesizing models. It is important not to miss components that are essential to the regulation mechanism in order to be able to explain the observations, but also not to consider components that do not play any role and hence penalize the construction and understanding of the system. If the interaction graph contains components that are not involved in the regulation of the observed biological behavior, many different functions can be attributed to these components without impacting the compatibility of the Boolean network with the observations. These components without importance for the behavior then strongly increase the number of solutions without bringing any information. Conversely, if no combination of functions can reproduce the data because key components and interactions of the process are missing, no model can be found. The complexity of this task currently limits the use of modeling.

With BoNesis, we propose a way to assist in the design of a relevant interaction graph with regard to observations. It offers to select, among a large interaction graph (as it can be extracted from a public interaction database such as DoRothEA (34) or Signor (43)), the components that can be included in a model to explain the observations.

To this end, we set two optimization criteria. Firstly, we want the solver to search for a compatible boolean network that maximizes the number of components in the models. An optimal solution is then a biggest boolean network, composed of components coming from the large original interaction graph, that can reproduce the observations. Yet, we also want the solution to maximize the number of components of a particular type, called *strong constants*. A strong constant is a component to which a constant function is assigned, and whose value remains constant to reproduce the observations within the dynamics of the Boolean network. Thus, within a Boolean network compatible with biological data, a node *A* is a strong constant if and only if *f* (*A*) = *v* with *v* ∈ {−1, 1} and that within the dynamics of the Boolean network it is possible to reproduce the data with the node *A* always equal to *v* (all the configurations *x* associated with the observations have *x*_*A*_ = *v*). In other words, it is a component having neither activator nor inhibitor and whose value can remain unchanged without preventing the Boolean network from being compatible with the observations. The particularity of these strong constants is that they can be removed from the domain without impacting compatibility with data behavior. Once the domain of interactions has been reduced to components that can explain the dynamics of the observations and that are not strong constants, we can limit ourselves to the maximum strongly connected component of this graph, which is particularly interesting to focus on the interactions that regulate the observed process.

#### Model synthesis

##### Exhaustive enumeration

BoNesis searches for all the Boolean networks of the same size than the input interaction network and whose dynamics exactly reproduce those of the data (via the defined dynamical constraints). Hence, when BoNesis is used for model synthesis without optimisation criteria, all the Boolean networks output from BoNesis are models of the same relevance with respect to the dynamics to be modeled.

##### Partial enumeration with diversity

For biological applications, it is frequent that an exhaustive enumeration of models is not judicious given that biological observations are rarely constraining enough considering a large size interaction network. Indeed, it is sufficient that a few components have few dynamic information for having an explosion of the number of models.

#### Specifics for case study 1

We first performed component selection (see previous section) from the 1 001 genes prior knowledge influence graph and the dynamical constraints of figure 2C. We extracted the largest strongly connected component of the resulting graph, leading to selecting 39 genes and 137 regulations. Then, from this 39 genes influence graph, we performed diverse sampling of 1 000 Boolean networks fulfilling the dynamical properties of figure 2C.

#### Specifics for case study 2

As described in the main text, the dynamical properties consisted of (1) existence of trajectories, (2) stability properties, and (3) absence of trajectories. Moreover, we add prior knowledge markers of the two differentiated cellular types: `ADIPOQ, FABP4, CEBPA, LPL` for adipocytes, and `ALPL, HEY1, SP7` for osteoblasts.

In a first stage, we performed gene selection on the METACORE(r) prior knowledge influence graph by identifying Boolean networks that maximize, by decreasing priority, the inclusion of a prior knowledge markers, the number of genes whose dynamics can be explained, and the number of strong constants, that will be removed (see previous section). For this first staged, we ignored constraints on the absence of trajectory for complexity reasons. We performed this gene selection for each replicate and each binarization method.

We then analyzed the binary valued inferred by BoNesis for the selected genes that have not been classified by binarization methods in the stable phenotypes. We identified 19 genes (`ATF2, CLOCK, CTCF, CUX1, GATA6, NFYB, REL, RELA, SMAD1, SMAD2, SMAD4, STAT3, TGIF1, TRP53, TTF1, USF1, USF2, YY1, ZFP148`) that have been inferred to have different binary states, while the data show no clear statistical ground for supporting these different qualitative states. We performed again the gene selection stage with the additional constraint that the binary value of these 19 genes must be equal in all phenotypes. Depending on the binarization method and replicates, different sets of genes have been selected, with, in some cases, multiple optimal solutions. Then, for each optimal solution, we extracted the sub-GRN consisting of the identified non-constant genes, and performed diverse sampling of Boolean networks taking into account the whole dynamical constraints, including on the absence of trajectories.

### Boolean network clustering and complexity analysis (Case study 1)

#### Complexity of the models per clusters

To explore the three highlighted groups, we study if the complexity of the functions of a model is characteristic of the group to which the model belongs (figures S4 and S5). Specifically, figure S4 shows, for each group, the percentage of the functions of its models according to the number of clauses constituting the functions. figure S5 shows, for each group, the percentage of clauses constituting its model functions according to the number of components in the clauses. We observe that red models are significantly less complex than those of the other two groups, with functions composed of a smaller number of clauses, themselves composed of a smaller number of components compared to the orange and green group models.

#### Complexity of the invariable functions of the models per clusters

Among the functions constituting the models, some are invariable between models of the same group beyond the 12 invariable functions which are common to all groups. We therefore have three sets of invariable functions that seem to indicate three key patterns of interactions to reproduce the data dynamics. Figure S6 shows, for each group, the percentage of its invariable functions according to their number of clauses. Again, a clear difference in complexity between the red group and the others is highlighted. All the functions shared by the red models have a single clause, whereas it represents for green and orange models respectively less than half and one third of the functions (some of their invariable functions having even respectively up to 7 and 8 clauses). Hence, the red group models appear to be more parsimonious explanations of the regulation mechanism of hematopoiesis than the two other groups.

### Computational prediction of reprogramming determinants (case study 2)

The prediction of the combinations of perturbations for the case study 2 has been performed via the software CABEAN (77). CABEAN implements several methods for the source-target control of asynchronous Boolean networks, and these methods can be used to identify the minimal and exact control sets of perturbations that ensure the inevitable reachability of the target attractor from a given source attractor. Based on the application time of the perturbations, CABEAN supports several types of controls: instantaneous control applies perturbations instantaneously; temporary control applies perturbations for sufficient time and then releases them to retrieve the original dynamics; and permanent control applies the control for all the following time steps. All these control methods (80) are based on the computation of strong and weak basins of attractors, which explores both the structure and dynamical properties of asynchronous Boolean networks. More specifically, an instantaneous control drives the network dynamics from the source attractor to a state in the strong basin of the target attractor, from which there only exist paths to the target attractor. On the other hand, temporary control and permanent control can make use of the spontaneous evolutions of the network dynamics by moving into first the weak basin of the target attractor, from which there exist paths to the target attractor. To ensure the inevitable reachability of the target attractor, a temporary control should drive the network dynamics to a state in the strong basin of the target attractor at the end of control, while a permanent control stirs the network from the source state to a state in the strong basin of the target attractor in the resulting transition system under control. More recently, CABEAN (78) has been extended with target control methods for asynchronous Boolean networks (79), which can be used to identify perturbations that can drive the dynamics of a Boolean network from any initial state to the desired target attractor.

In this way, CABEAN fits well with the objective for case study 2, i.e., enforcing the convergence to `OD15` state from `AD15` state. CABEAN was applied to each inferred Boolean networks from BoNesis to compute combinations of temporary perturbations (gene knockout), and it identified combinations of up to 5 simultaneous perturbations leading to a reprogramming from `AD15` to OD15. We kept combinations of perturbations that have been identified in at least 10% of the individual models given to CABEAN. In the end, we have obtained 34 different combinations of 2 to 4 simultaneous perturbations, which are given as input for the next analysis steps.

### Stochastic simulations of ensembles of Boolean networks (case study 2)

To perform simulations on our ensembles of models, we developed a new version of MaBoSS, the stochastic Boolean simulator(76), called Ensemble-MaBoSS(11). MaBoSS usually takes a single Boolean model and performs a large number of simulations to obtain many stochastic trajectories. It later uses this set of trajectories to compute time-dependent probabilities for every visited Boolean state. During these simulations, it also stores every fixed point observed and produces the set of fixed points observed in all these simulations. To perform the simulation of an ensemble of models, we need to compute an equal set of trajectories for every model of the ensemble. We can then use the consolidated set of trajectories from all the models, and compute the time-dependent probabilities specific to the whole ensemble of models. We can also produce the same list To analyze the composition of the ensemble, we also compute the time-dependent probabilities for each model. This way, we get individual results, that can then be used for ensemble analysis such as model clustering.

### Wetlab experiments (case study 2)

#### Cell culture

The mouse bone marrow-derived stroma cell line ST2 was established from Whitlock-Witte type long-term bone marrow culture of BC8 mice (65). The ST2 cells were cultured in growth medium: Roswell Park Memorial Institute (RPMI) 1640 medium (Gibco, 32404-014) supplemented with 10% fetal bovine serum (FBS) (Gibco, 10270-106) and 1% L-Glutamine (Lonza, BE17-605E) at 37°C, 5% CO2. All experiments were carried out with cells less than 10 passages. For differentiation of adipocytes, ST2 cells were seeded 2 days before differentiation. The cells reached 100% confluency after 24 hours of culture, and were further maintained in growth medium for 24 hours (Day 0). Adipogenic differentiation was initiated on Day 0 by culturing in adipogenic differentiation medium I consisting of growth medium, 0.5 mM isobutylmethylxanthine (IBMX) (Sigma-Aldrich, I5879), 0.25 *μ*M dexamethasone (DEXA) (Sigma-Aldrich, D4902) and 5 *μ*g/mL insulin (Sigma-Aldrich, I9278). From Day 2, the cells were cultured in adipogenic differentiation medium II consisting of growth medium, 500 nM rosiglitazone (RGZ) (Sigma-Aldrich, R2408) and 5 *μ*g/mL insulin (Sigma-Aldrich, I9278). The adipogenic differentiation medium II was replaced every second day.

For differentiation of osteoblasts, ST2 cells were seeded 2 days before differentiation, and reached 100% confluency after 24 hours of culture, and were further maintained in growth medium for 24 hours (Day 0). Osteogenic differentiation was initiated on Day 0 by culturing in osteogenic differentiation medium consisting of growth medium and 100 ng/mL bone morphogenetic protein-4 (BMP4) (PeproTech, 315-27). The osteogenic differentiation medium was replaced every second day.

#### RNA extraction and cDNA synthesis

Total RNA was extracted from cells using Quick-RNATM MiniPrep (Zymo Research, R1055). RNA concentration was measured by Nanodrop 2000c (Thermo Fisher Scientific, E597). cDNA was synthesized with the following cocktail: 1 *μ*g total RNA, 0.5 mM dNTPs (Thermo Fisher Scientific, R0181), 2.5 *μ*M oligo dT-primer (Eurofins GmbH), 1 U/*μ*L Ribolock RNase inhibitor (Thermo Fisher Scientific, EO0381), and 1 U/*μ*L RevertAid Reverse transcriptase (Thermo Fisher Scientific, EP0352). The cocktail was maintained at 42^°^C for 1 hour, following 70^°^C for 10 min to stop the reaction.

#### Reverse transcription real-time quantitative PCR (RT-qPCR)

RT-qPCR was performed to measure the RNA expression using the Applied Biosystems 7500 Fast Real-Time PCR System. Each reaction contained the following cocktail: 5 *μ*L of cDNA, 5 *μ*L of primer mix (forward and reverse primers, both in 2*μ*M concentration), and 10 *μ*L of Absolute Blue qPCR SYBR Green Low ROX Mix (Thermo Fisher Scientific, AB4322B). The PCR reaction were the following: 95^°^C for 15 min and repeating 40 cycles of 95^°^C for 15 s, 55^°^C for 15 s, following 72^°^C for 30 s. The gene expression level was calculated using the 2-(ΔΔCt) method (48). The ΔΔCt refers to (ΔCt(target gene) – ΔCt(housekeeping gene)) from the treatment (ΔCt(target gene) – ΔCt(housekeeping gene)) from the control. Rpl13a was used as the housekeeping gene and the primer sequences can be found in the Supplementary Table 2.

#### Viral transduction

Day 9 adipocytes were transduced with lentivirus (Sirion Biotech Gmbh; details can be found in the Supplementary Table 2). For a well of 48-well plate, the estimated cell number was 200 000. The amount of lentivirus was calculated to achieve expected multiplicity of infection (MOI). The transduction cocktail was as followed: lentivirus stock diluted with RPKM and 8 *μ*g/mL polybrene transfection reagent (EMD Millipore, TR-1003-G) to achieve a final volume of 100 *μ*L. The culture medium was removed, and the cells were washed once with 1xDPBS (Gibco, 14190-094), then supplemented with the transduction cocktail. The cells were kept in the transduction cocktail for 6 hours. After the 6-hour transduction, the transduction cocktail was removed, and the cells were supplemented with the growth medium. The transduction efficiency was controlled by observed GFP and RFP levels by microscopy.

#### RNA interference

On Day 1 post-transduction (Day 1 PT), the cells were transfected with siRNA according to manufacturer’s recommendation (Horizon Discovery; details can be found in the Supplementary Table 2). In brief, siRNA was diluted to 5 *μ*M solution in DNase/RNase free water (Invitrogen, 10977-035). In separate tubes, siRNA and DharmaFECT 1 transfection reagent (Horizon Discovery, T-2001-03) were diluted with RPMI. To prepare the transfection cocktail for 1 well of 48-well plate, in Tube 1, 25 *μ*L of the siRNA in serum-free medium was prepared by adding 1.25 *μ*L of 5 *μ*M siRNA to 23.75 *μ*L of RPMI. In Tube 2, 1 *μ*L of the DharmaFECT 1 in serum-free medium was prepared by adding 1 *μ*L of DharmaFECT 1 to 24 *μ*L of RPMI. The mixture was incubated for 5 min, then two tubes were mixed following incubation of 20 min. After incubation, 200 *μ*L of growth medium was added to achieve a final volume of 250 *μ*L. The culture medium was removed, and the cells were supplemented with the transfection medium for 24 hours. For single and double siRNA transfection, the final concentration was maintained at 100 nM.

#### Single cell RNA-seq

The single cell RNA-seq (scRNA-seq) was performed according to Chrominum Next GEM Single Cell 3’ Reagent Kits v3.1 Review D (CG000204). The ChromiumTM Next GEM Single Cell 3’ GEM, Library Gel Bead Kit 3.1 (1000128) consisted the following: Chrominum Next GEM Single Cell 3’ Gel Bead Kit v3.1, 4 (1000129), Chromium Next GEM Single Cell 3’ GEM Kit v3.1 (1000130), Chromium Next GEM Single Cell 3’ Library Kit v3.1 (1000158), Single Index Kit T SetA (1000213), DynabeadsTM MyOneTM SILANE (2000048). To achieve single cell suspension, the cells were first treated with 1 mg/mL Collagenase A (Roche, 10103586001) for 15 min. After Collagenase A treatment, the cells were trypsinised with 150 *μ*L Trypsin (Lonza, BE17-161E) for 5 min. The trypsin was quenched with 500 *μ*L growth medium. Wide-bore tips (Thermo Scientific ART 1000G Self-Sealing Barrier Pipet Tips, 2079G) were used to pipet up and down. The suspension was centrifuged for 1000 rpm for 5 min, and the supernatant was removed. Single cell suspension was achieved by adding 400 *μ*L growth medium and pipetted up and down with wide-bore tips. The cell number was determined with C-CHIP (NanoEntek, DHC-N01). The suspension was centrifuged again for 1000 rpm for 5 min, and the supernatant was removed and replaced with 1xDPBS + 0.04% BSA to achieve the target cell number. The suspension was filtered through 40 *μ*M Flowmi Cell strainer (Merck, BAH136800040-50EA). The cell number was determined again with C-CHIP, which would be used to determine the input for scRNA-seq. In brief, cells was loaded with a targeted recovery rate of 10 000 cells per sample. scRNA-seq library quality were assessed by Agilent DNA High sensitivity Bioanalyzer chip (Agilent, 5067-4626) and further sequenced on a 150 cycles High Output Kit using Illumina NextSeqTM 500 with targeted sequencing depth of 20 000 read pairs per cell.

## Data and code availability

The software BoNesis, employed for Boolean network inference, is available at https://bnediction.github.io/bonesis under the GPLv3-compatible free software license CeCiLL. The software Cabean, employed for reprogramming prediction, is available at https://satoss.uni.lu/software/CABEAN. Both tool are available through the CoLoMoTo Docker distribution (61) at https://colomoto.github.io/colomoto-docker/ with persistently archived Docker images. *Case study 1:* data and scripts are available at https://github.com/StephanieChevalier/notebooks_for_bonesis/tree/main/applications/hematopoiesis. *Case study 2:* Matlab code for statistical analysis and binarization of bulk RNA-seq data is available at https://github.com/sysbiolux/algorecell; data and scripts for Boolean network inference and control predictions are are available at at https://sdrive.cnrs.fr/s/nrB72LpBspeKHEp.

## SUPPLEMENTARY FILES

- Supplementary Table 1 (SupplTable1.xlsx). Differential gene expression analysis from scRNA-seq RE1 experiment.
- Supplementary Table 2 (SupplTable2.xlsx). Primer sequences, siRNAs, and viral constructs used in the study. The primers, siRNAs, and lentiviral constructs are detailed in the indicated work sheets, respectively.

**Figure S1:**
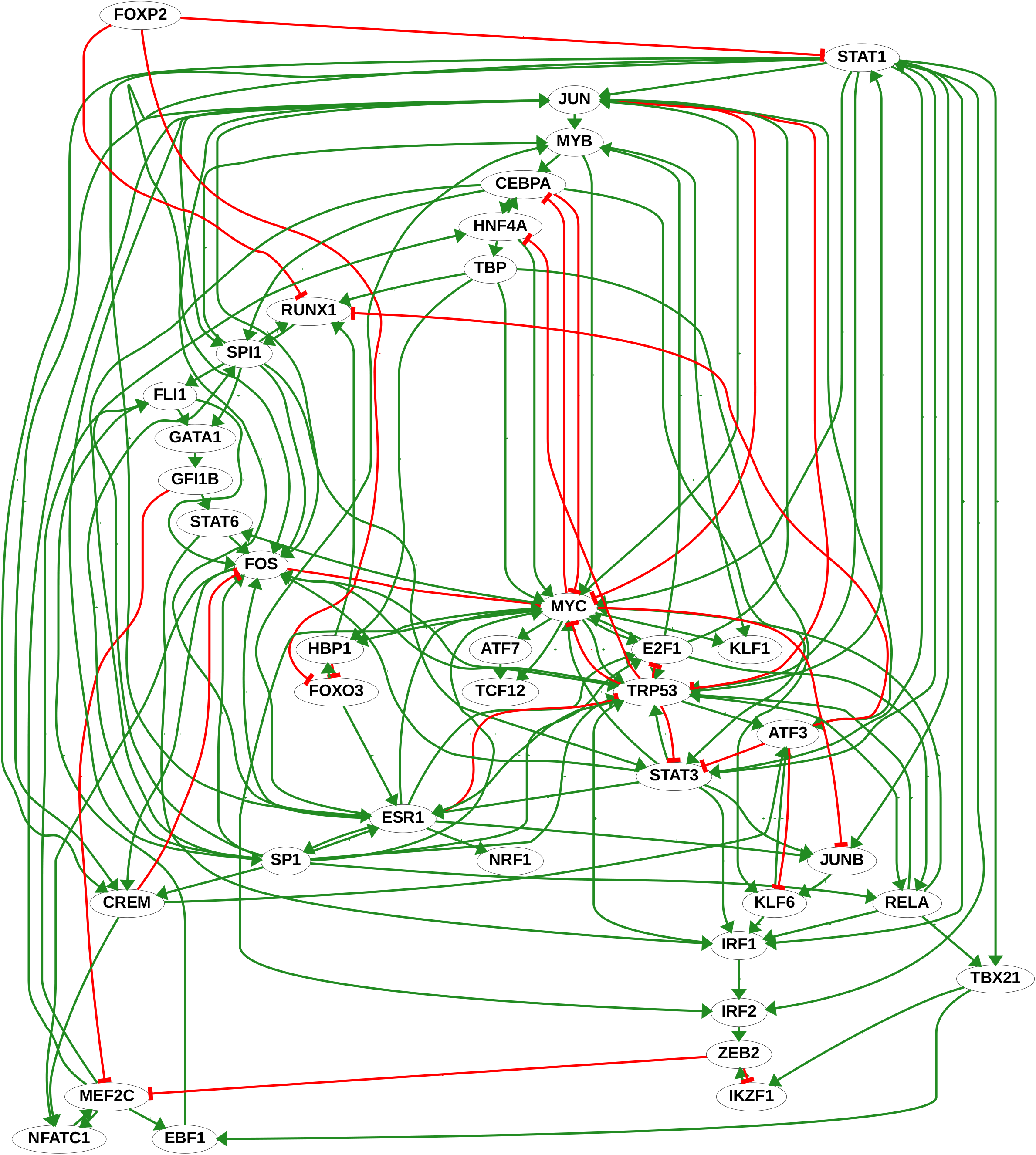
Network of 39 components and 137 arcs obtained by component selection using BoNesis.

**Figure S2:**
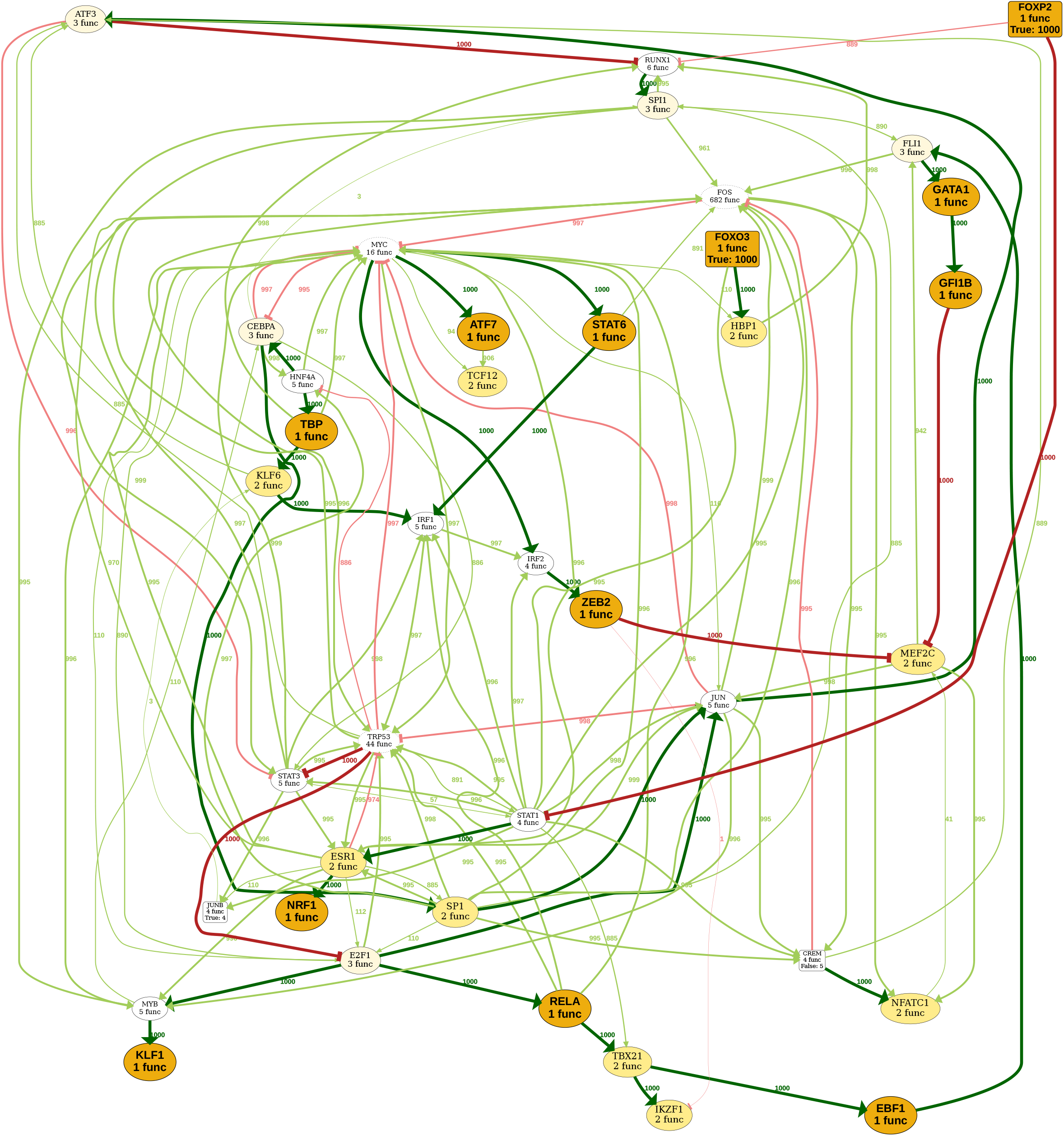
Variability analysis Boolean functions in sampled ensemble of models. The label of each node includes the number of different local Boolean functions in the sampled Boolean network set. Orange indicates a unique shared function, yellow only 2 different functions. Edge edge is labeled with the number of Boolean networks that utilizes the influence (over the 1000 sampled), with its thickness scaled accordingly.

**Figure S3:**
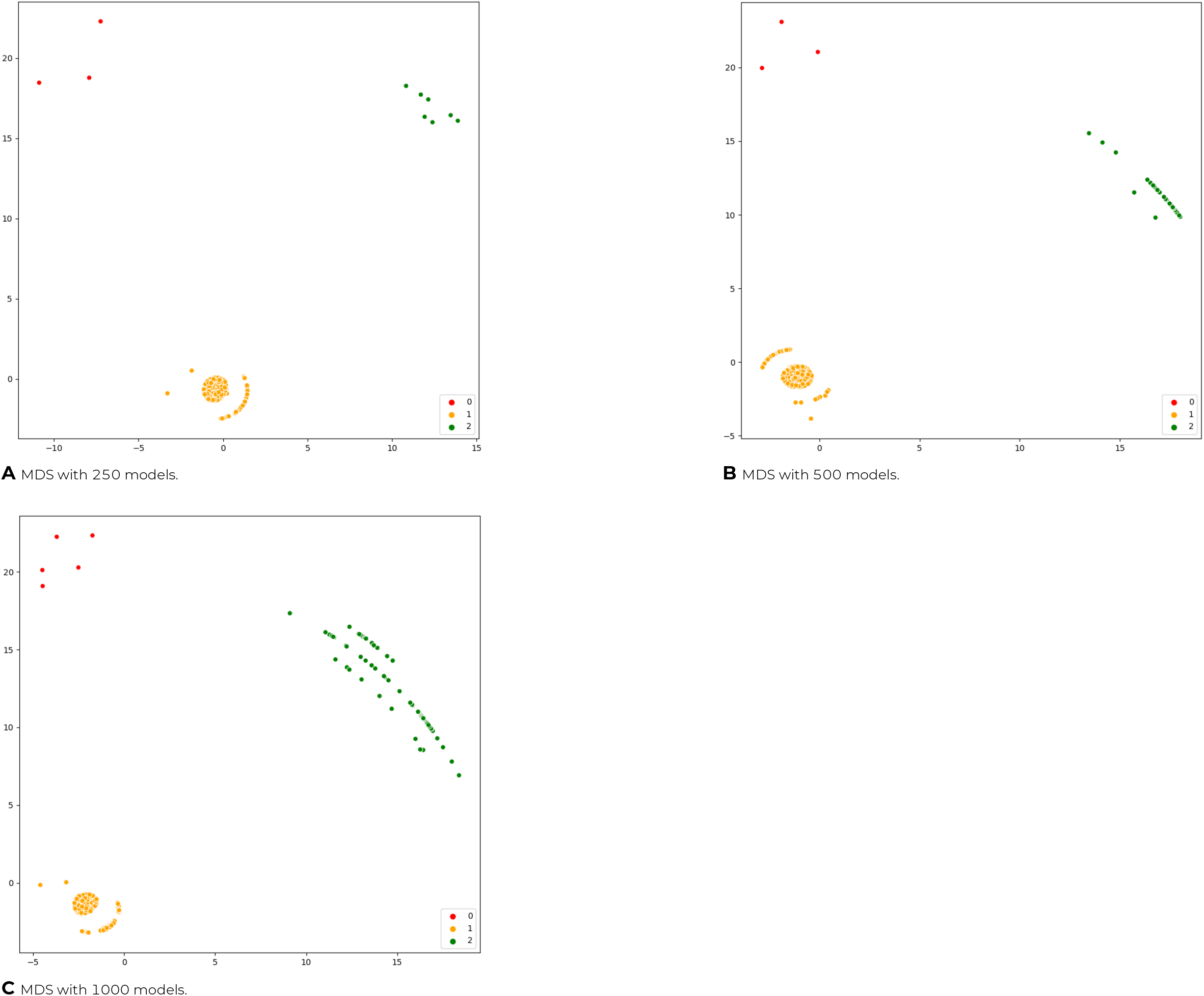
MDS highlights 3 groups of models.

**Figure S4:**
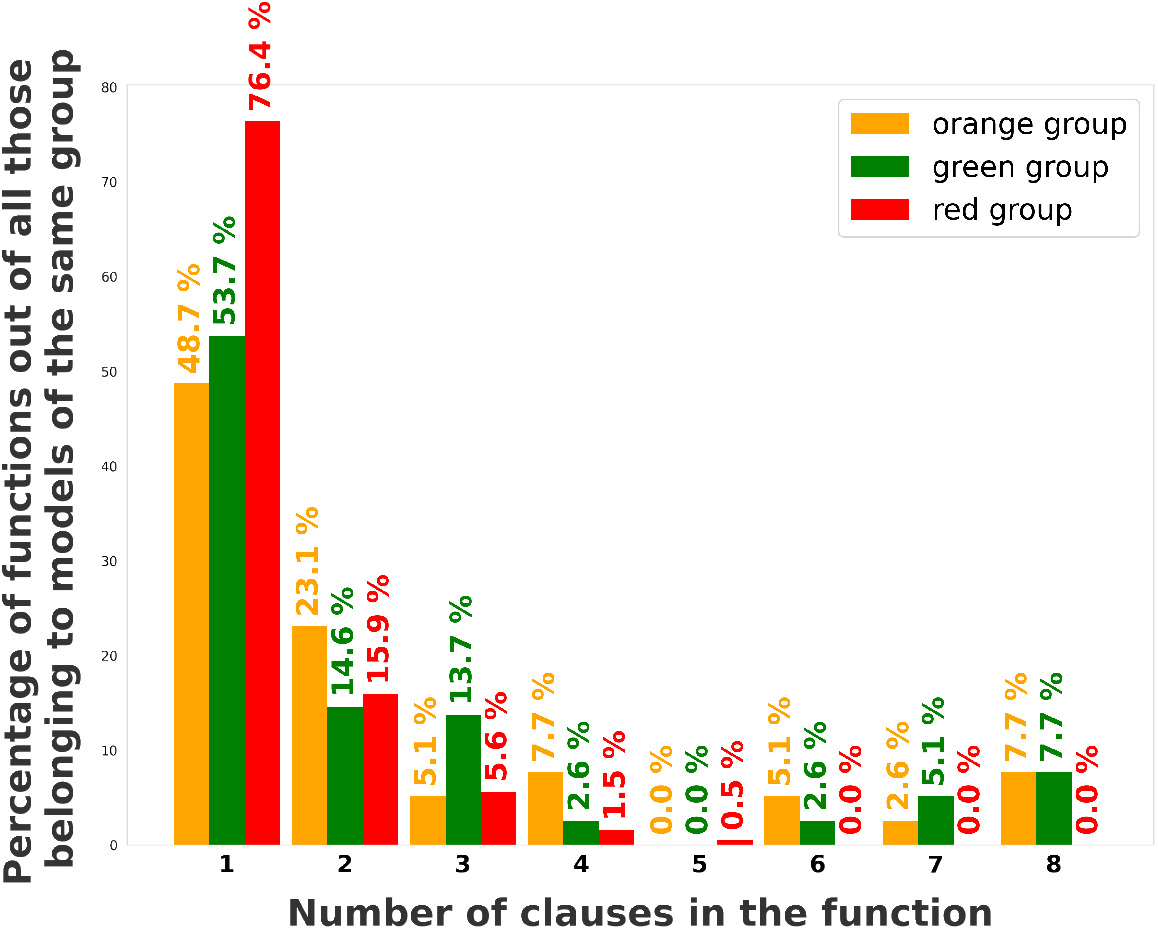
Distribution of the functions of the models according to the number of clauses they are made up, by group.

**Figure S5:**
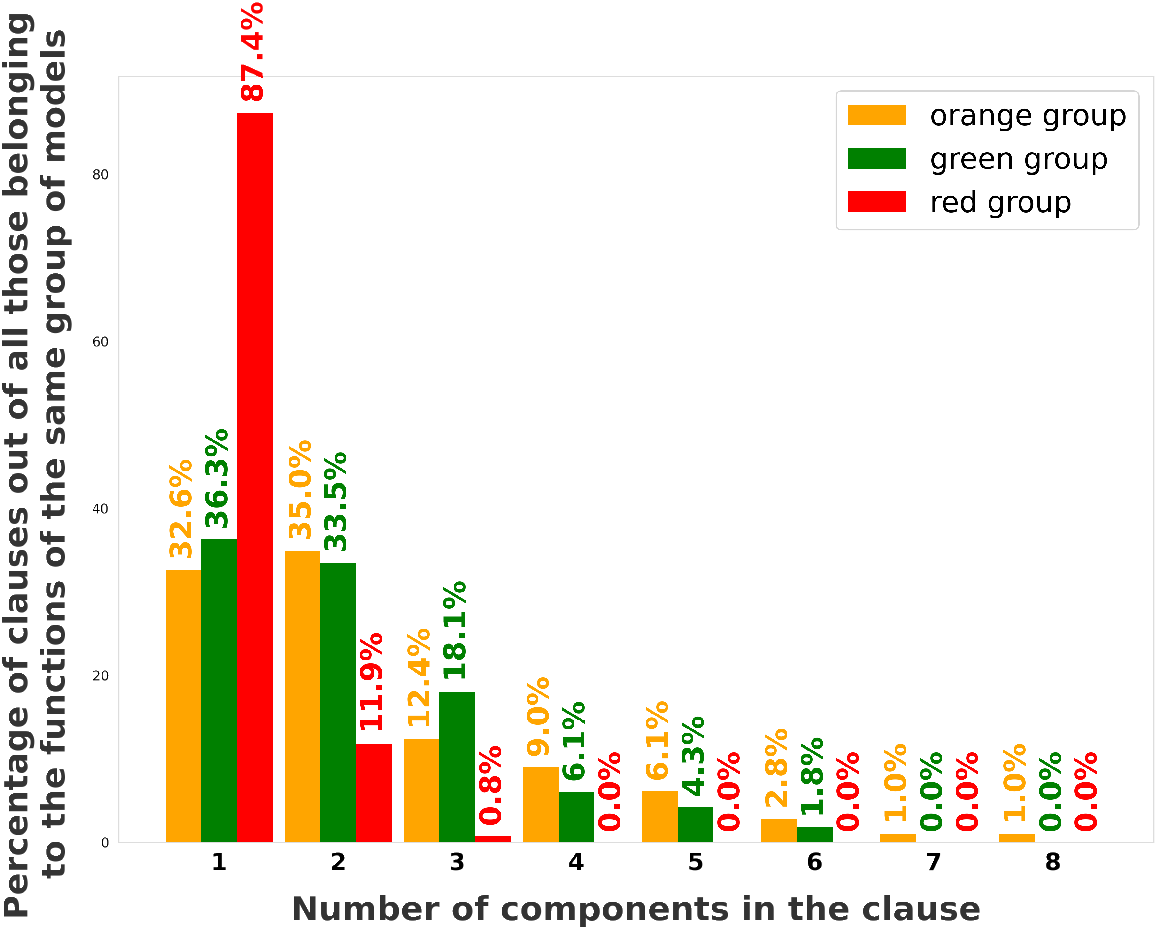
Distribution of the clauses of the models according to the number of components they are made up, by group.

**Figure S6:**
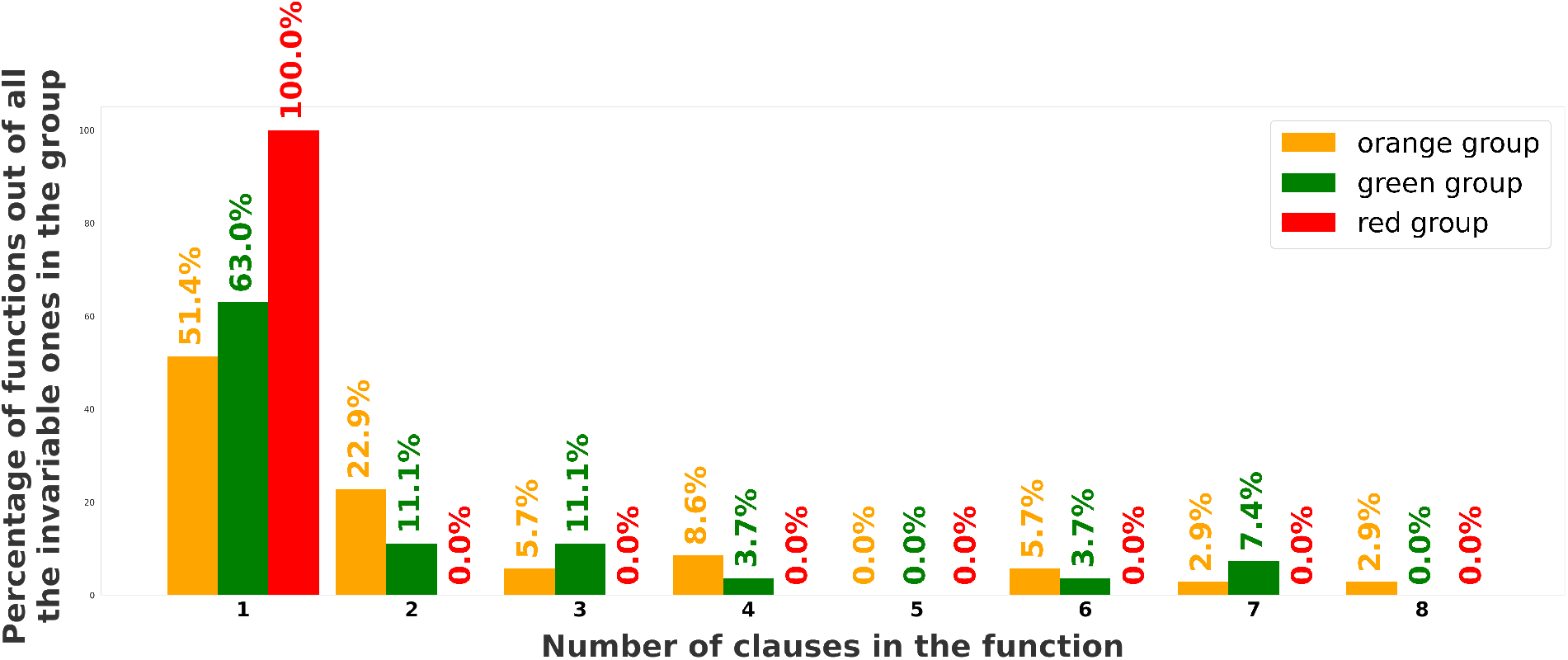
Distribution of the invariable functions of the models according to the number of clauses they are made up, by group.

**Figure S7:**
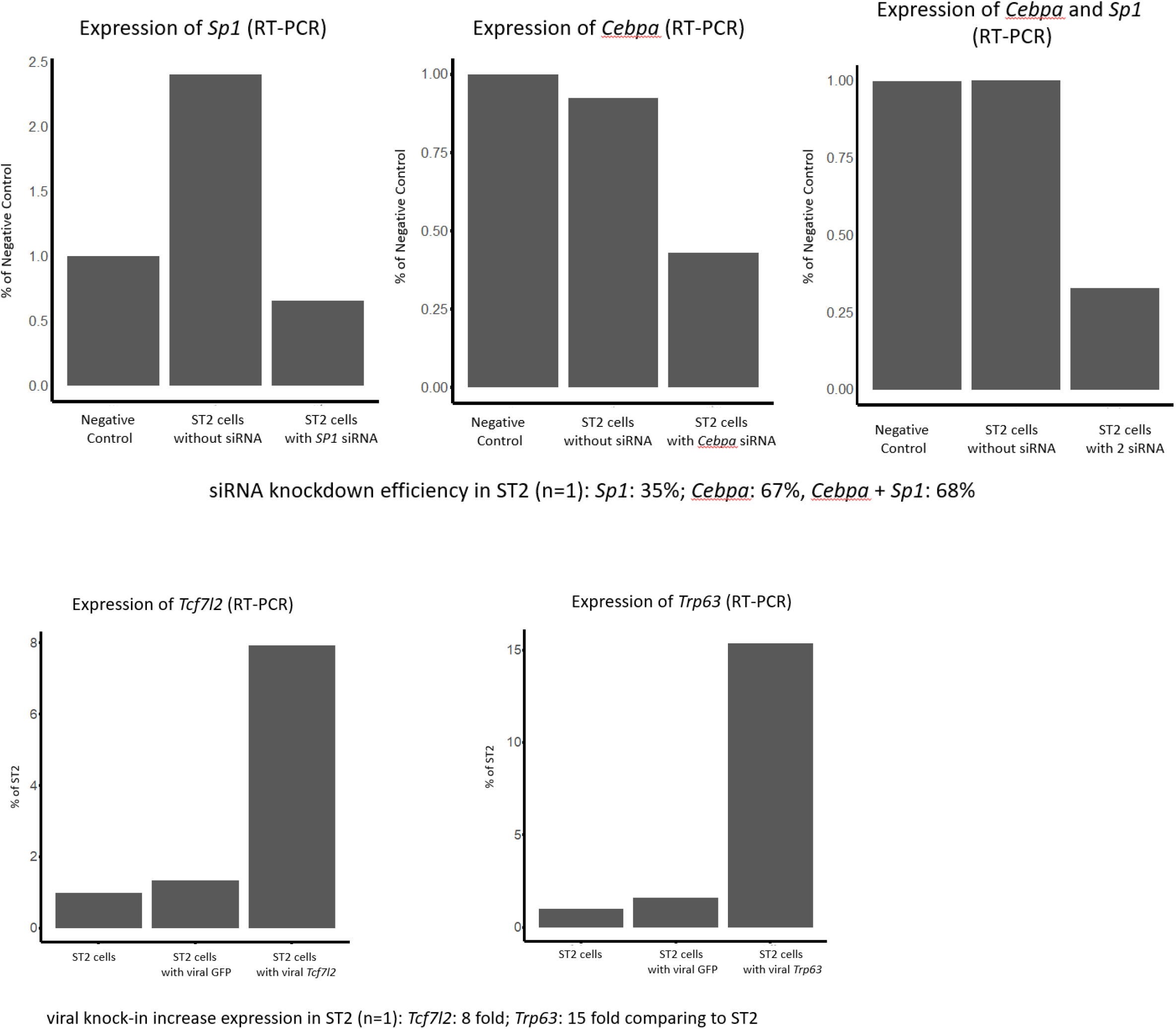
Experimental validation of case study 2: siRNA and viral transfection test

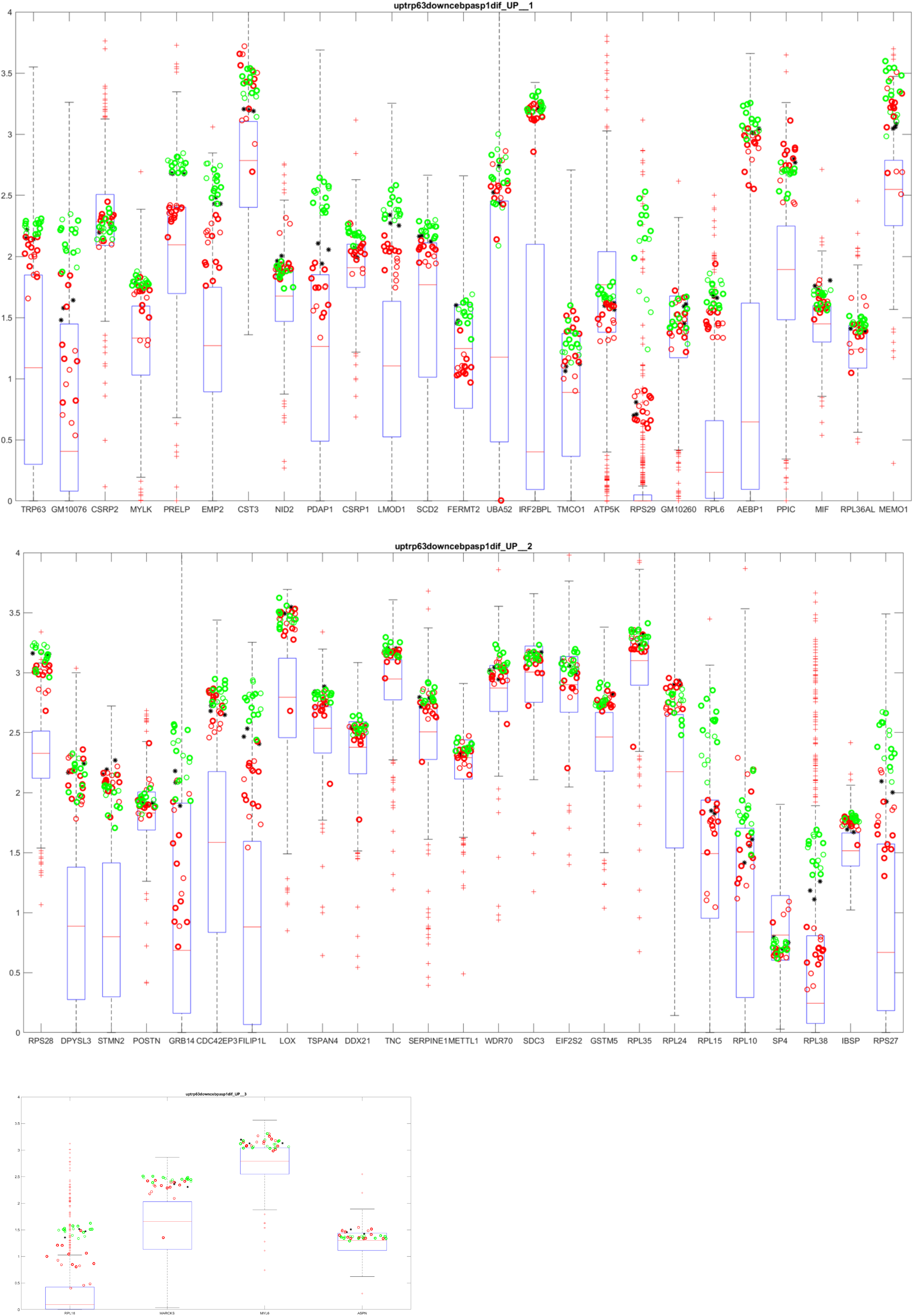

**Figure S9:**
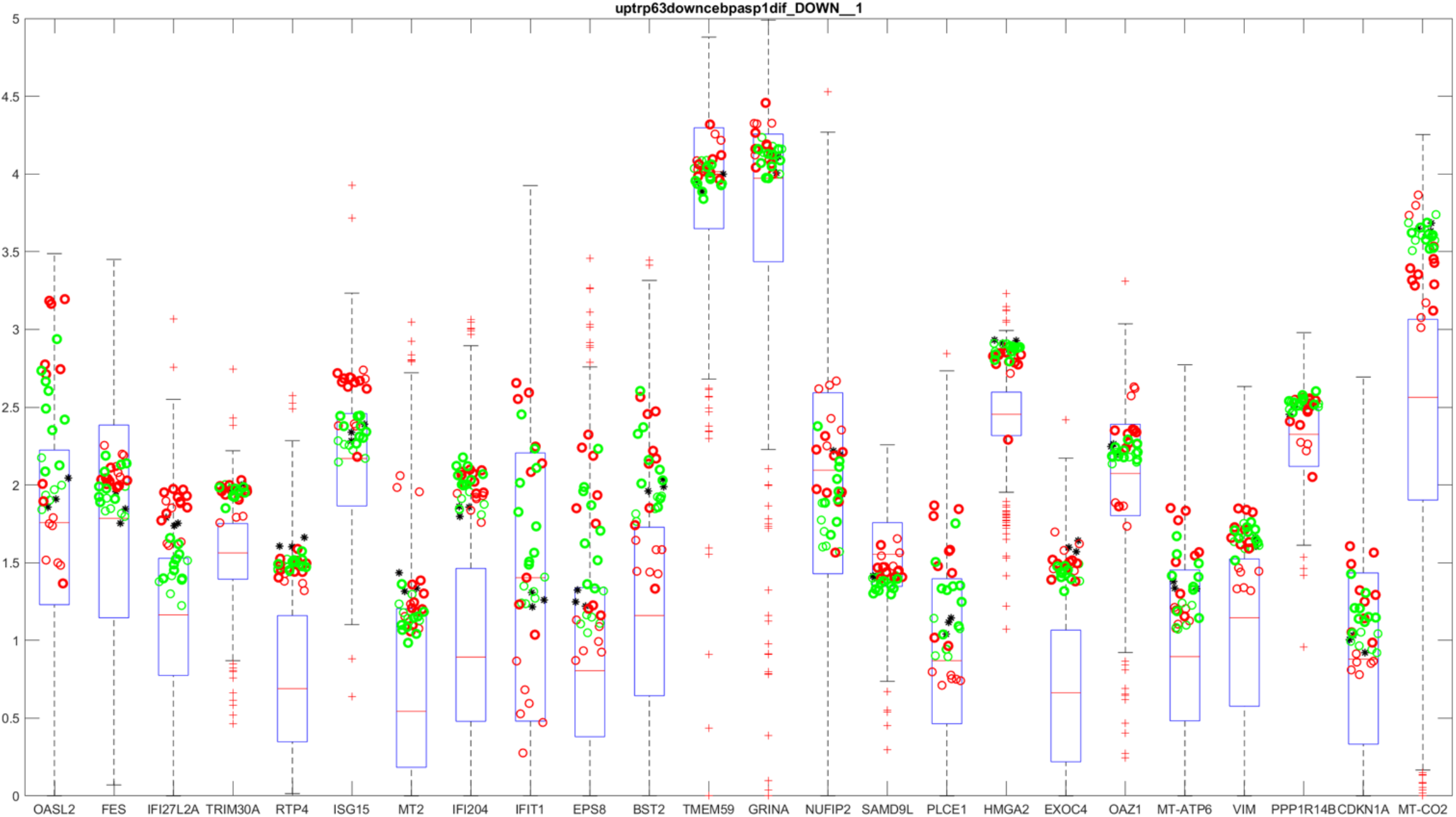
Differentially expressed downregulated genes in the RE1 single cell experiment are partially also downregulated in the bulk RNA-seq data of osteoblasts vs adipocytes. Bulk RNA-seq data (TPM) of osteoblasts (green) and adipocytes (red) are plotted for downregulated sc DEGs (x-axis). Boxplots indicate the gene specific backgrounds from 63 mouse tissues (see methods).

